# Bamboozle: A bioinformatic tool for identification and quantification of intraspecific barcodes

**DOI:** 10.1101/2023.03.16.532925

**Authors:** Matthew I M Pinder, Björn Andersson, Karin Rengefors, Hannah Blossom, Marie Svensson, Mats Töpel

## Abstract

Evolutionary changes in populations of microbes, such as microalgae, cannot be traced using conventional metabarcoding loci as they lack intraspecific resolution. Consequently, selection and competition processes amongst strains of the same species cannot be resolved without elaborate isolation, culturing, and genotyping efforts. Bamboozle, a new bioinformatic tool introduced here, scans a species’ entire genome and identifies allele-rich barcodes that enable direct identification of different strains from a common population, and a single DNA sample, using amplicon sequencing. We demonstrate its usefulness by identifying hypervariable barcoding loci (<500 bp) from genomic data in two microalgal species, the diploid diatom *Skeletonema marinoi*, and the haploid chlorophyte *Chlamydomonas reinhardtii*. Across the genomes, only 26 loci capable of resolving all available strains’ genotypes were identified, all of which are within protein-coding genes of variable metabolic function. Single nucleotide polymorphisms (SNPs) provided the most reliable genetic markers, and amongst 55 strains of *S. marinoi,* three 500 bp loci contained, on average, 46 SNPs, 103 unique alleles, and displayed 100% heterozygosity. The prevalence of heterozygosity was identified as a novel opportunity to improve strain quantification and detect false positive artefacts during denoising of amplicon sequences. Finally, we illustrate how metabarcoding of a single genetic locus can be used to track strain abundances of 58 strains of *S. marinoi* in an artificial selection experiment. As future genomics datasets become available and DNA sequencing technologies develop, Bamboozle has flexible user settings enabling optimal barcodes to be designed for other species and applications.

## Introduction

Microalgae are a diverse paraphyletic group of aquatic microbes responsible for half of global primary production (Falkowski et al. 1998; Field et al. 1998). Many species have enormous population sizes compared with multicellular organisms and contain some of the highest intraspecific genetic diversity observed within eukaryotes (Flowers et al. 2015). Despite their ecological importance, knowledge about individual species’ functional diversity and evolutionary potential is limited (Godhe and Rynearson 2017). Microalgal blooms can contain thousands to millions of different clones (Sassenhagen et al. 2021), but designs of evolution experiments are currently constrained to only one or a few strains as they generally utilise clonal cultures, e.g., (Lohbeck et al. 2012; Schaum et al. 2017; Schaum et al. 2018; Sefbom et al. 2015; Wolf et al. 2019). If these strains are co-cultured in competition or *in situ*, as needed for many evolutionary and ecological questions, the individual strains cannot readily be identified or quantified using existing microscopic or genotyping technologies. Currently, genotyping of microalgae relies heavily on microsatellites (Rengefors et al. 2017), although newer sequencing methods such as RAD-seq provide a method for fine-scale genotyping and genomic information (Rengefors et al. 2021). Although suitable for population genetic and genomic studies, these methods still rely on single DNA samples from each strain for genotype identification. This severely restricts these methods’ usefulness in quantitative evolutionary studies, where strain isolation is time-consuming or even impossible if the species is difficult to culture (Rengefors et al. 2021), or if the population’s clonal diversity is large (Schaum et al. 2017; Scheinin et al. 2015).

An alternative approach to track cellular abundances involves using massively parallel amplicon sequencing of a diverse locus. The main advantage of such an approach is that millions of individual DNA molecules, and by extension the cells they originate from, can be quantified from a single DNA sample. This simple principle - metabarcoding - has revolutionised the field of microbial ecology in the past decade (Hebert et al. 2003; Taberlet et al. 2012). Depending on the taxonomic group and the resolution required, various highly conserved genes are commonly used in metabarcoding, including ribosomal RNA genes (16S, 18S, 23S, or 28S), mitochondrial cytochrome c-oxidase subunit 1 (COI), and the chloroplastic ribulose-1,5-bisphosphate carboxylase/oxygenase large subunit (*rbcL)* (Guo et al. 2015; Pujari et al. 2019; Yamada et al. 2017). Metabarcoding enables a relatively simple and standardised quantification of ecosystem-wide parameters, such as diversity, presence of low abundance or cryptic taxa, and community composition. However, the conserved nature of the available metabarcoding loci restricts their usefulness as intraspecific markers (Canesi and Rynearson 2016; Godhe et al. 2006; Guo et al. 2015).

With a sufficiently diverse intraspecific barcode marker it should be possible to track the abundances and quantify fitness of individual strains from a mixed microbial population. Such an approach would be analogous to ‘barcoding analysis by sequencing’, or Bar-seq experiments, where unique barcodes are used to replace genes in knock-out mutants (Robinson et al. 2014). Bar-seq experiments then uses amplicon sequencing to quantify fitness changes in thousands of non-lethal gene deletion mutants simultaneously, using a co-culture experimental design (Kim et al. 2010; Li et al. 2019; Robinson et al. 2014; Smith et al. 2009), and it has accelerated the functional annotations of several microbial species’ genomes (Dent et al. 2005; Giaever and Nislow 2014; Han et al. 2010). Access to ‘intraspecific’ DNA barcode loci in microalgae and other microbes would provide a powerful tool to address fundamental questions about their evolution and population structures, analogous to how metabarcoding and Bar-seq have radically changed our understanding of microbial ecology and phenotypic functions of genes in the last decade.

Genome sequencing projects are beginning to resolve the genetic diversity between species of microalgae (Armbrust et al. 2004; Bowler et al. 2008; Merchant et al. 2007; Mock et al. 2017; Read et al. 2013; Worden et al. 2009), and population genomic datasets are becoming increasingly available (Blanc-Mathieu et al. 2017; Flowers et al. 2015; Osuna-Cruz et al. 2020; Rastogi et al. 2020; von Dassow et al. 2015), providing information about intraspecific genetic diversity. From this data, it should be possible to identify novel barcoding loci with intraspecific resolution. Here we introduce Bamboozle, a bioinformatic approach that uses whole genome sequencing data to identify intraspecific barcodes by scanning a species’ entire genome for the most suitable sites. As a proof-of-concept, we identify and design primers for a set of barcodes in the diploid marine diatom *Skeletonema marinoi* and the haploid model chlorophyte *Chlamydomonas reinhardtii*. Furthermore, we demonstrate how one single barcode locus can be used to track strain selection in an artificial evolution experiment incorporating 58 environmental strains of *S. marinoi* from two Baltic Sea inlets on Sweden’s east coast. Based on the data from this experiment we identify opportunities, as well as challenges, in employing amplicon sequencing to address questions regarding evolutionary processes in microbes. Although we see obvious applications in microalgae and other microbes, Bamboozle can identify intraspecific barcodes for both haploid and diploid species when provided with a reference genome and whole genome sequencing data from multiple individuals.

### New approaches

Our aim with the Bamboozle tool was to enable identification of novel intraspecific barcoding loci with resolution down to the level of genotypes originating from one common population. To this end, we needed to identify loci fulfilling three criteria that render them suitable as barcoding loci, as outlined by Kress and Erickson (2008): 1) they should be short enough to allow amplification and sequencing with current technologies, 2) they should have significant interspecific (in our case intraspecific) variability, and 3) they should have conserved flanking regions that allow the binding of primers. Our approach builds on existing approaches for identifying novel barcoding loci by scanning the entire genome (Angers-Loustau et al. 2016; Paracchini et al. 2017). In contrast to the aforementioned studies, we focus on identification of barcodes with intraspecific resolution, implement several filters to optimise allele richness, exclude unsuitable genomic regions, and minimise the amount of manual screening needed.

The steps of the Bamboozle approach for barcode identification are outlined in Fig. 1 and described in detail below. The required input data includes a reference genome for the organism of interest, and whole genome sequencing (WGS) data from the strains to be analysed. Throughout the paper we use the strain concept (Lakeman et al. 2009), which for most practical purposes is synonymous with genotype in multicellular organisms. The WGS data is required in two formats, one pair per strain: 1) a sorted BAM file generated by aligning the WGS reads to the reference, and 2) single nucleotide polymorphism (SNP) data in VCF format generated from the aforementioned BAM file using GATK (Van der Auwera and O’Connor 2020).

**Figure 1:**
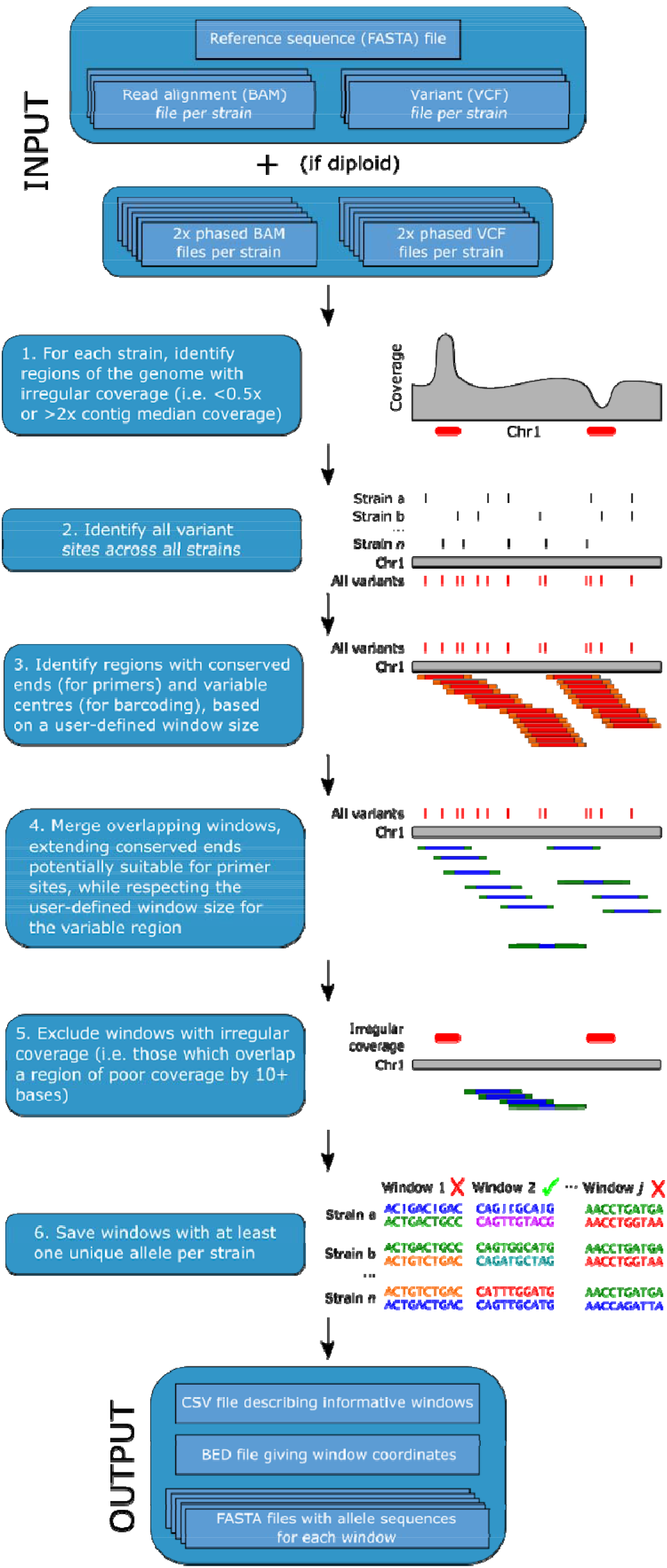
Schematic summarising the steps of the Bamboozle pipeline, as well as input and output files. Each step is described in more detail in the New Approaches section.

In the case of diploid species, phased BAM and VCF files (i.e., where data from each allele is in a separate file) are also required to analyse the sequences of individual alleles. The main Bamboozle tool, written and tested in Python version 3.7.10, consists of six main steps, detailed below.

1. *Identification of regions of unusual coverage.* When mapping whole genome shotgun sequencing reads to a reference genome, one expects a similar read coverage across the entirety of the reference. While short regions deviating from this expectation could result from potentially informative deletions or insertions in the mapped strain versus the reference, longer such regions could be due to misassemblies in the reference genome, repeat regions with significant length differences between strains, or gene duplications. As these phenomena could be problematic in identifying appropriate barcoding loci and downstream amplicon sequencing, SAMtools (Danecek et al. 2021) and Bedtools (Quinlan and Hall 2010) are called by Bamboozle and used to determine the median read depth of each contig in each strain (`samtools depth`), and to identify regions which deviate heavily from this median value (`bedtools genomecov`). By default, regions with coverage less than 50% or more than 200% of the contig median in that strain are flagged, in order to filter out large heterozygous deletions and gene duplications, respectively.
2. *Identification of variant positions.* Using a custom algorithm written in Python version 3.7.10, the locations of all variants in each strain are extracted from their respective VCF files and consolidated.
3. *Genome screening based on conserved sites.* As one of the core requirements for a suitable barcode is conserved flanking regions for primer binding, windows containing such conserved regions are identified in this step. Bamboozle traverses each contig/chromosome (step size 1; window size defined by the user), saving for further analysis those windows with conserved regions (i.e., regions without any SNP’s) at each end, based on the variant list from step 2. The size of the window and the conserved end regions can be defined by the user with the `--window_size` and `--primer_size` options, respectively, to account for different amplification and sequencing strategies.
4. *Merging of overlapping windows.* Where both conserved end regions of two windows overlap, these two are merged. For example, a 500 bp window with conserved regions from contig positions 1-21 and 480-500, and a second such window with conserved regions from contig positions 2-22 and 481-501, would be merged to form a single 501 bp window with conserved regions from contig positions 1-22 and 480-501. This is done to reduce the number of overlapping windows to be screened manually after Bamboozle outputs the end results. For a visual example, see step 4 of Fig. 1.
5. *Exclusion of windows with unusual read coverage.* The merged windows are now compared to the coverage data generated in step 1, and windows that overlap with a region of read coverage fluctuation by ten or more bases are excluded. As length variations can be potentially valuable in terms of differentiating between strains, allowing short (<10 bp) stretches of irregular coverage (indicative of short indel regions) is intended as a compromise between the potential for informative windows and the complications such regions may introduce, as noted in step 1.
6. *Identifying informative loci.* As a unique sequence is required to identify and quantify each strain in, for example, a strain selection or phenotyping experiment, SAMtools (`samtools faidx`) and BCFtools (`bcftools consensus`) (Danecek et al. 2021) are applied to each remaining window to generate consensus sequences for each strain (and each allele, in the case of a diploid organism) using the reference genome and the respective VCF files. The individual allele sequences are then analysed to determine if each strain in the dataset contains at least one unique allele, in which case the window is reported to the user as a potentially suitable barcoding locus.

As output, Bamboozle reports the identified barcoding loci in 1) a tab-separated file, giving coordinates and metadata for each locus, 2) a BED file of locus coordinates, intended for easy visualisation, such as in a genome browser, and 3) a multi-FASTA file for each locus, containing the sequence of each allele in each strain. This enables quick and easy manual quality control of the output, and design of primers for PCR tests and amplicon sequencing. As a proof-of-concept, we performed amplicon sequencing on one locus identified in the diploid *S. marinoi* and used a custom-made script that merges amplicons from paired-end Illumina data using BBMerge (Bushnell et al. 2017), denoises the data using DADA2 (Callahan et al. 2016), and takes advantage of heterozygosity to quality control the output and translate allele abundances into counts of individual strains. Bamboozle offers a complete pipeline from WGS data through to amplicon sequencing analysis of the barcoding loci, and its performance has been fine-tuned and benchmarked using the diploid diatom *S. marinoi*.

## Results

### Performance of the bioinformatic pipeline: millions of potential barcoding loci reduced to twenty-six

As expected, common metabarcoding loci lacked intraspecific resolution in *S. marinoi*, with 80-100% of strains having the same single allele (Table S1). We applied Bamboozle to WGS data from 54 strains of *S. marinoi*, to identify novel barcoding loci for use with 2×301 bp paired-end Illumina MiSeq sequencing. Two of the initial 56 datasets were excluded from the analysis, one due to low WGS read coverage, and one for being a clone of another strain in the study. Initial attempts to identify suitable barcoding loci of 300-400 bp failed in this species (data not shown), so we scanned for loci of 500 bp. Within the *S. marinoi* genome, most (99.5%) of the 54,753,029 possible 500 bp windows were filtered out due to either coverage fluctuations (main problem, Table 1) or unconserved flanking regions. Of the remaining 263,000 windows, the average frequency of SNP positions was 12±11, and the average number of alleles was 14 (Table 1). Four potential barcoding loci were identified as containing at least one unique allele per strain and passing all filters (Table 1). Three of these, plus an additional hypervariable locus (*Sm_C2W24*) from a previous iteration of Bamboozle that did not include a coverage filter (step 1, Fig. 1), were selected for further evaluation. Compared to other 500 bp windows passing the read depth and conserved flanking region filters, three barcodes proved especially rich in terms of SNPs and unique alleles, falling in the 97.6th to 99.9th percentiles, respectively (Fig. 2A and B). *Sm_C2W24* was predicted to contain an even higher SNP frequency (Fig. 2B). The *Sm_C2W24* locus was retained through the analysis to assess the performance of Bamboozle in identifying loci without coverage filter implementation.

**Figure 2:**
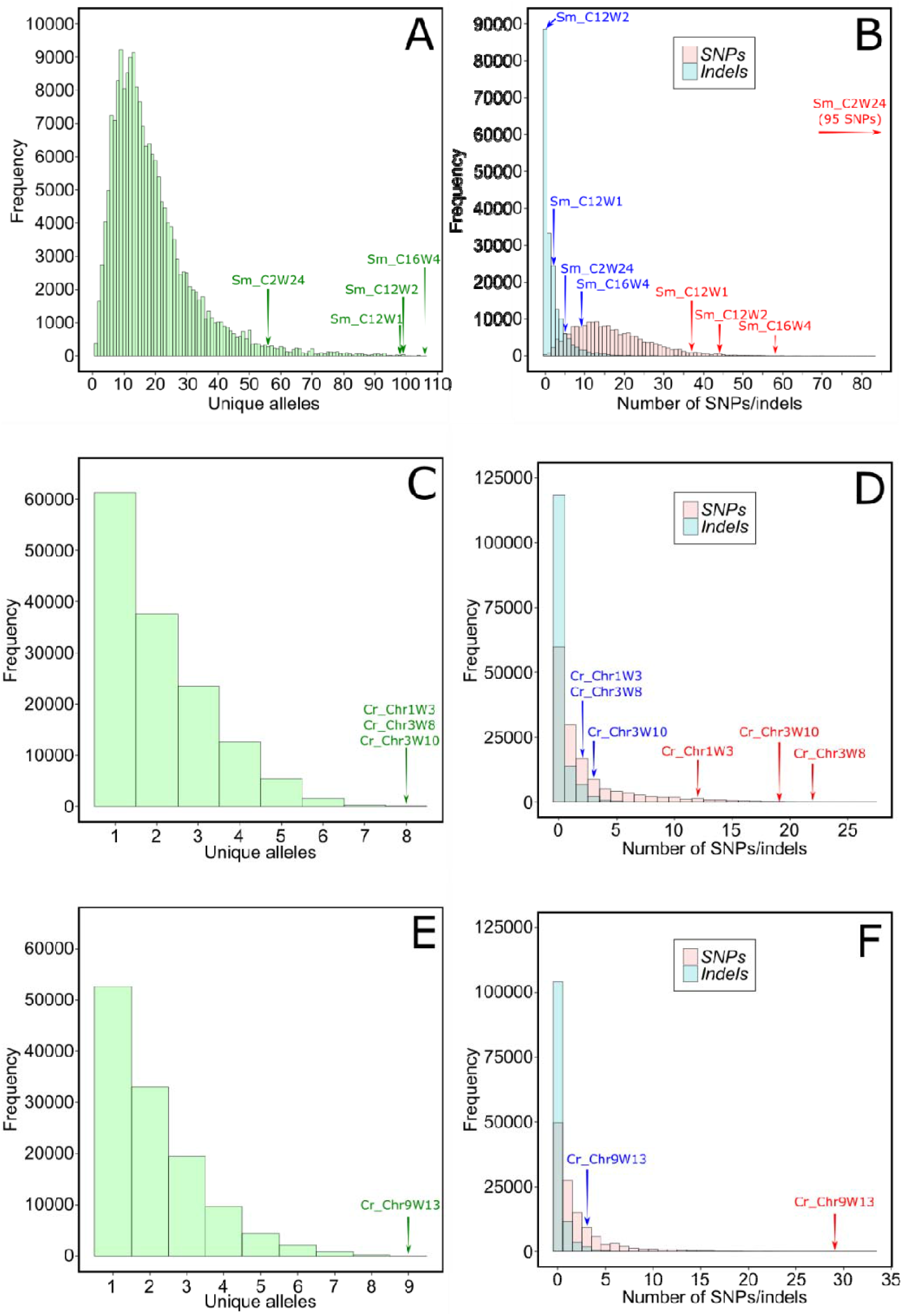
Histograms showing the predicted number of unique alleles (panels A, C, and E), and predicted frequency of SNPs and indels (panels B, D, and F), at each window of 500 bp (*S. marinoi*) or 300 bp (*C. reinhardtii*). Shown are data from all genomic windows that passed Bamboozle’s filters for coverage depth and flanking region conservation, as described in the New Approaches section. In the case of *S. marinoi*, only nuclear contigs are included. Panels A and B depict figures from *S. marinoi*; panels C and D depict figures from *C. reinhardtii* mating type +; panels E and F depict figures from *C. reinhardtii* mating type -. Arrows indicate the relative locations of the selected barcoding loci in terms of predicted SNP (red) and indel (blue) frequency, and unique alleles (green). Note that *Sm_C2W24* was predicted with an earlier version of Bamboozle, resulting in deviant values compared to other loci.

**Table 1.**
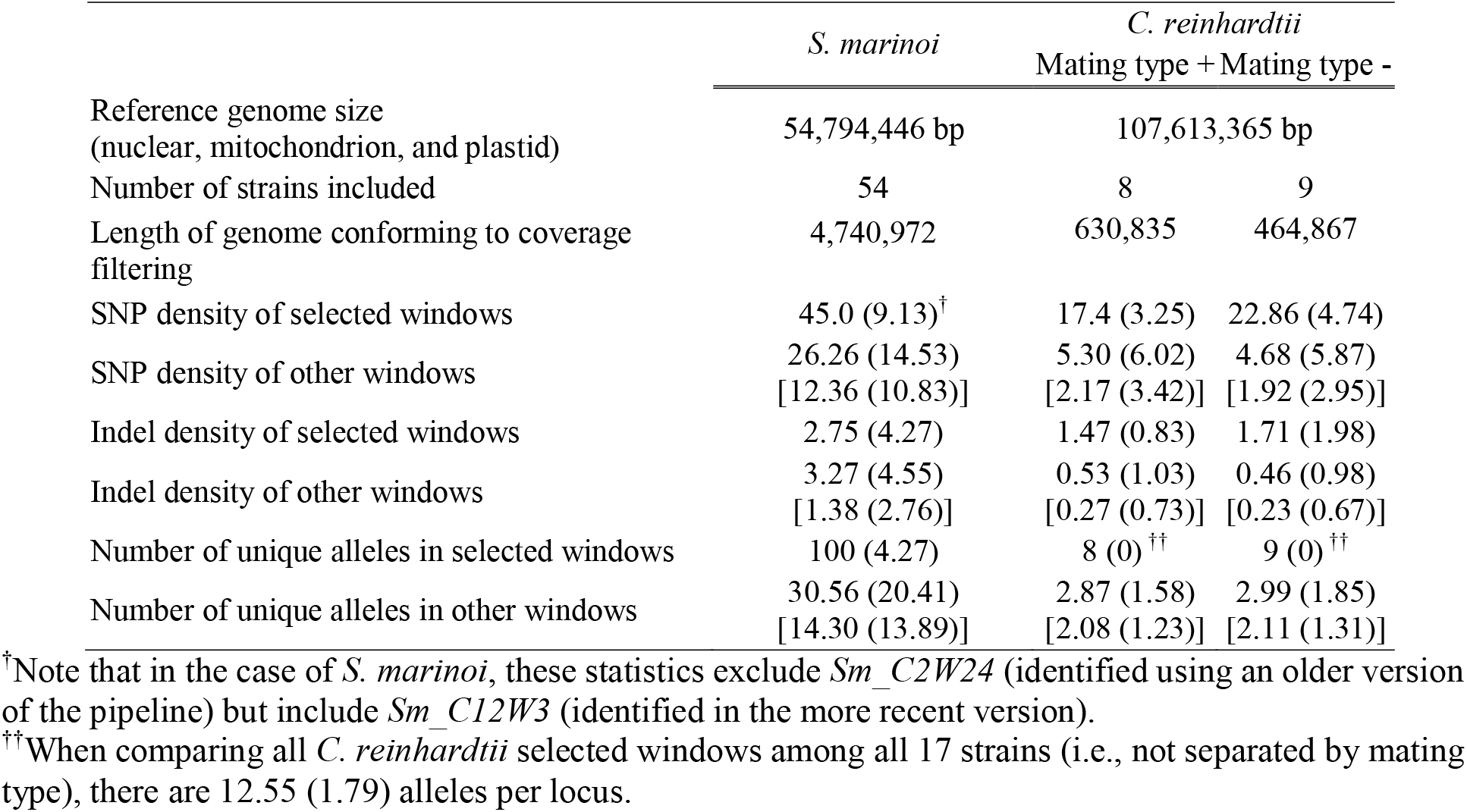
Statistical parameters from Bamboozles genomic scan for barcode windows. Shown are average (standard deviation) for the variable barcoding loci identified by Bamboozle (selected windows), with regard to the number of predicted SNPs, indels, and unique alleles per locus, in contrast to average statistics for the genome as a whole, obtained using a sliding-window approach (other windows). Where two sets of values are given, the first gives statistics are for those windows where only the coverage filter is applied, and the second (in square brackets) gives statistics for those windows were both the coverage filter and the conserved end region filter are applied.

The Bamboozle tool was also applied to seventeen haploid *C. reinhardtii* strains to identify loci of ∼300 bp. While the initial analysis on all strains returned no suitable barcoding loci, separation by mating type (eight mt+ strains and nine mt-strains) was successful. From a reference genome of 107,613,365 bp [nuclear genome, mitochondrion, plastid, and mt- locus (Merchant et al. 2007)], with 107,161,335 potential 300 bp windows, only 137,000 loci (0.13%) contained less than two-fold coverage deviation compared with the contig median (again the main issue, Table 1) and had conserved flanking regions. About 40% of remaining windows contained only one allele in the population, and less than 1% contained more than seven (Fig. 2C and E). Only 22 of these loci were identified by Bamboozle as having one unique allele for all strain genomes - fifteen for mt+ and seven for mt- (Table S2). In addition to being significantly more diverse in allelic richness, these Bamboozle-identified loci also contained a much higher number of SNPs than other loci across the genome (three out of four ranked in the 99.8th percentile; Fig. 2D and F).

### In silico annotation of intraspecific barcodes and common features

All 26 barcodes identified in *S. marinoi* and *C. reinhardtii* were inside predicted protein-coding genes (Table S2). Several barcodes were also in close physical proximity within the same, or adjacent genes, e.g., *Sm_C12W1*, *-2*, and *-3*, located within a 2,197 bp region containing two gene models (Sm_t00009768-RA and Sm_t00009769-RA), as well as *Cr_Chr9W-12* and *-13* inside *CHLRE_09g389134v5*, and *Cr_Chr11W-16, -17,* and *-18,* inside *CHLRE_11g467528v5*. The predicted function of the proteins did not have a consistent pattern across the two species, and ranged from ribosomal proteins (Sm_t00008465-RA [ribosome biogenesis protein WDR12] and *CHLRE_10g447800v5* [60kDa SS-A/Ro ribonucleoprotein]), to enzymes (*CHLRE_03g207250v5* [putative glutamine synthetase] and *CHLRE_13g592050v5* [allantoinase]) and ion transporters (*CHLRE_11g467528v5* [calcium channel]). Eleven barcodes were located inside genes without conserved domains or with conserved Domains of Unknown Function (DUF).

Four barcodes of each species were selected for primer design and further barcode development (Fig. 3). Among these, no part of the ∼500 bp barcoding loci was intronic in *S. marinoi*. Instead, three out of four barcodes spanned domains with repetitive amino acid regions (*Sm_C2W24*, *Sm_C12W1*, and *Sm_C16W4*) with the conserved primer sites anchored inside the same or flanking conserved domains (Fig. 3A). In contrast, for *C. reinhardtii,* the major part of the 300 bp barcoding loci was intronic, while the conserved primer sites were located either in introns or exons (Fig. 3C). A more detailed annotation of the barcode-containing genes is provided in the Supplemental Information.

**Figure 3:**
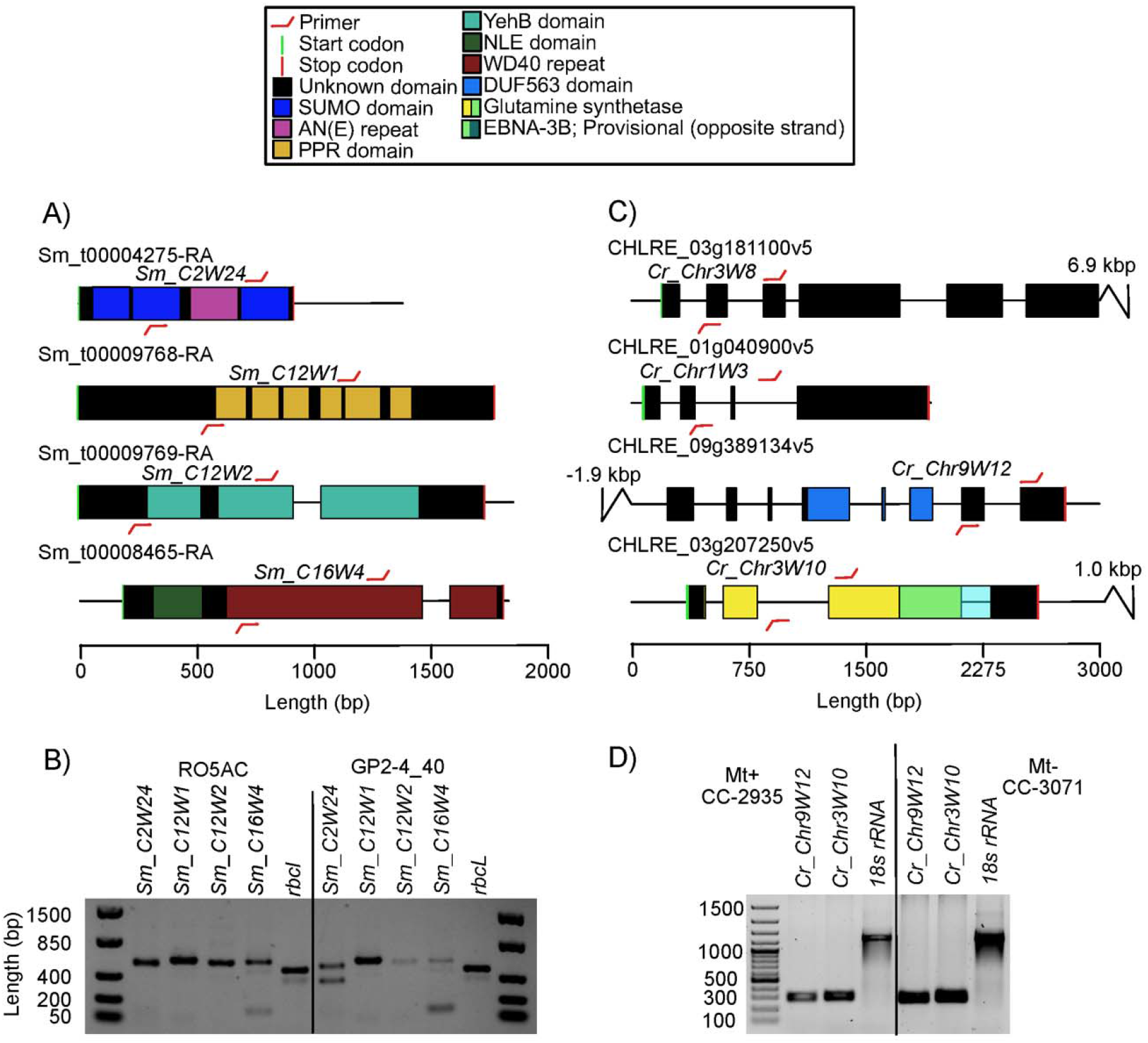
Genomic location of the four barcoding loci identified in *S. marinoi* and selected four loci in *C. reinhardtii,* corresponding to those that primers were designed and evaluated for (for complete list of all loci, and brief gene model annotations, see Table S2). A and C) Gene models with functional domains, primer sites, and variable regions in *S. marinoi* and *C. reinhardtii,* respectively. B and D) Gel electrophoresis of PCR products in two strains of *S. marinoi* and one strain each of mating type + and - in *C. reinhardtii,* of loci with good amplification results. Abbreviations – SUMO: Small Ubiquitin-like Modifier; AN(E): the three amino acids Alanine, Asparagine, and Glutamic acid [occasionally replaced by Glycine]; PPR: pentatricopeptide [RNA binding]; YehB: Uncharacterised membrane protein [partial, UPF0754 family]; NLE: NUC135 domain located in N terminal of WD40 repeats; WD40: domain involved in various cellular function including signal transduction, pre-mRNA processing and cytoskeleton assembly; DUF563: Domain of Unknown Function; EBNA-3B: PHA03378 superfamily domain (located on the reverse strand). Glutamine synthetase is represented by its PLN02284 superfamily domain and overlap with EBNA-3B is shown in green. Note that the primer symbols are not drawn to scale but the tip of each symbol indicates the exact location where they end and the variable region begins. Department of Biology, Lund University, Lund, Sweden

As future barcode applications may involve co-cultivation with other species, or *in situ* experiments in natural microalgal communities, we evaluated to what extent the primers were species-specific. In the target species, primers could be designed to amplify the selected loci around the 21 bp conserved regions (Fig. 3B and D), although strain amplification appeared uneven between strains for *Sm_C12W2, Sm_C16W4, Cr_Chr3W1*, and *Cr_Chr1W3*, suggesting possible PCR amplification bias. Homologous genes for the *S. marinoi* barcode loci were identified in three other species of centric diatoms within the order *Thalassiosirales* (*Thalassiosira pseudonana* and *Thalassiosira oceanica,* and *Skeletonema subsalsum*) (Table S3). In the two *Thalassiosira* species, each locus’s primer sites contained mismatches in at least three positions. However, in *S. subsalsum* the *Sm_C12W2* locus contained only one mismatch, and this was also the only locus that yielded a PCR product using *S. subsalsum* DNA as template. None of the primer pairs amplified in eleven other Baltic Sea phytoplankton species (Fig. S1A and B).

Although the 54 strains of *S. marinoi* all came from a remote population of *S. marinoi* inside the Baltic Sea, the primer sites were conserved also in three strains from the American Atlantic Coast and the Mediterranean Sea (Table S3), suggesting that the barcodes will function outside the local Baltic Sea population. PCRs of the *C. reinhardtii* barcode loci *Cr_Chr9W12* and *Cr_Chr3W10* did not amplify in two other chlorophytes, including another *Chlamydomonas* sp. strain, suggesting they could also be species-specific (Fig. S1C).

### Accuracy of bioinformatic predictions and amplicon sequencing of barcodes in S. marinoi

The allele sequences predicted by Bamboozle were validated using amplicon sequencing of individual strains’ PCR products, or Sanger sequencing for the haploid *C. reinhardtii*. This was done for two reasons: firstly, to generate the accurate allele sequences in the strains and contrast this with Bamboozle’s predictions based on genomic shotgun sequences, and secondly, to assess what PCR and sequencing artefacts were prevalent.

Although *Sm_C2W24* was identified by an earlier version of Bamboozle that did not employ a coverage filter, we retained this locus throughout the analysis to illustrate issues that micro- or minisatellite-like loci can produce in the pipeline. Bamboozle predicted that *Sm_C2W24* contained 95 SNPs and five indels within our studied population, but manual inspection indicated that some strains had low sequencing coverage across the most variable region. Amplicon sequencing revealed length differences of 198 bp amongst alleles of *Sm_C2W24* in the *S. marinoi* population, caused by variable copy number (2 to 24 copies) of a 9 bp repeat encoding Alanine-Asparagine-Glutamic acid (AN(E)) (length difference also confirmed by gel-electrophoresis, Fig. 3B). The variant calling algorithm of GATK, and by extension Bamboozle, had misinterpreted these repeats as a high abundance of SNPs (Table 2). Consequently, almost all allele-sequences in the different strains were incorrectly predicted, and per-base accuracy across the locus was only 82.6%. Hence, as outlined in Fig. 1, adjustments were made to Bamboozle to filter out similar regions, and *Sm_C2W24,* together with 90-99% of the genomes (Table 1), was disregarded in subsequent analysis.

**Table 2:**
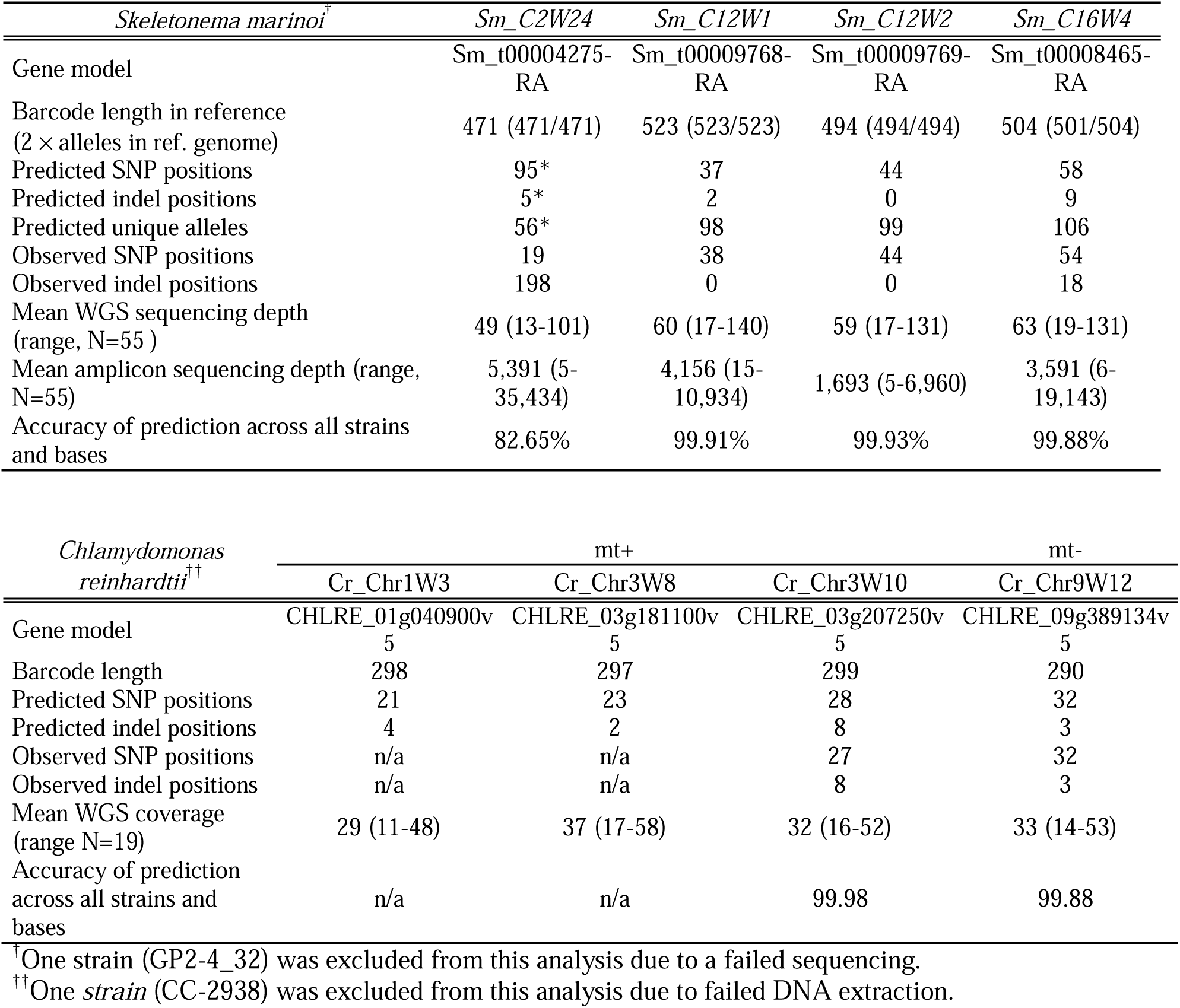
Accuracy of Bamboozle’s allele sequence predictions for selected barcodes in S. marinoi and C. reinhardtii. ‘Predictions’ corresponds to Bamboozle’s output sequences based on WGS data, and ‘observations’ are confirmations based on targeted PCR-based sequencing. C. reinhardtii loci with poor amplification and lack of Sanger sequencing data are shown as n/a. Sm_C2W24 results marked with * indicate results from an earlier iteration of the pipeline, which didn’t take coverage or allele phasing into account, and was variant-called using BCFtools (cf. GATK in the most recent iteration).

All strain-specific sequences of the three *S. marinoi* barcoding loci, as well as *Cr_Chr9W12* and *Cr_Chr3W10* in *C. reinhardtii*, identified with the coverage filter were predicted with >99.9% accuracy (Table 2). Looking into the reason for the remaining discrepancies, we discovered that some of the strains were triploid (GP2-4_54 and VG1-2_78; Fig. 4), confounding the software’s expectations of diploidy. In other cases, the inaccuracies were caused by insufficient filtering of the variant calling data, resulting in spurious base-calling errors.

**Figure 4:**
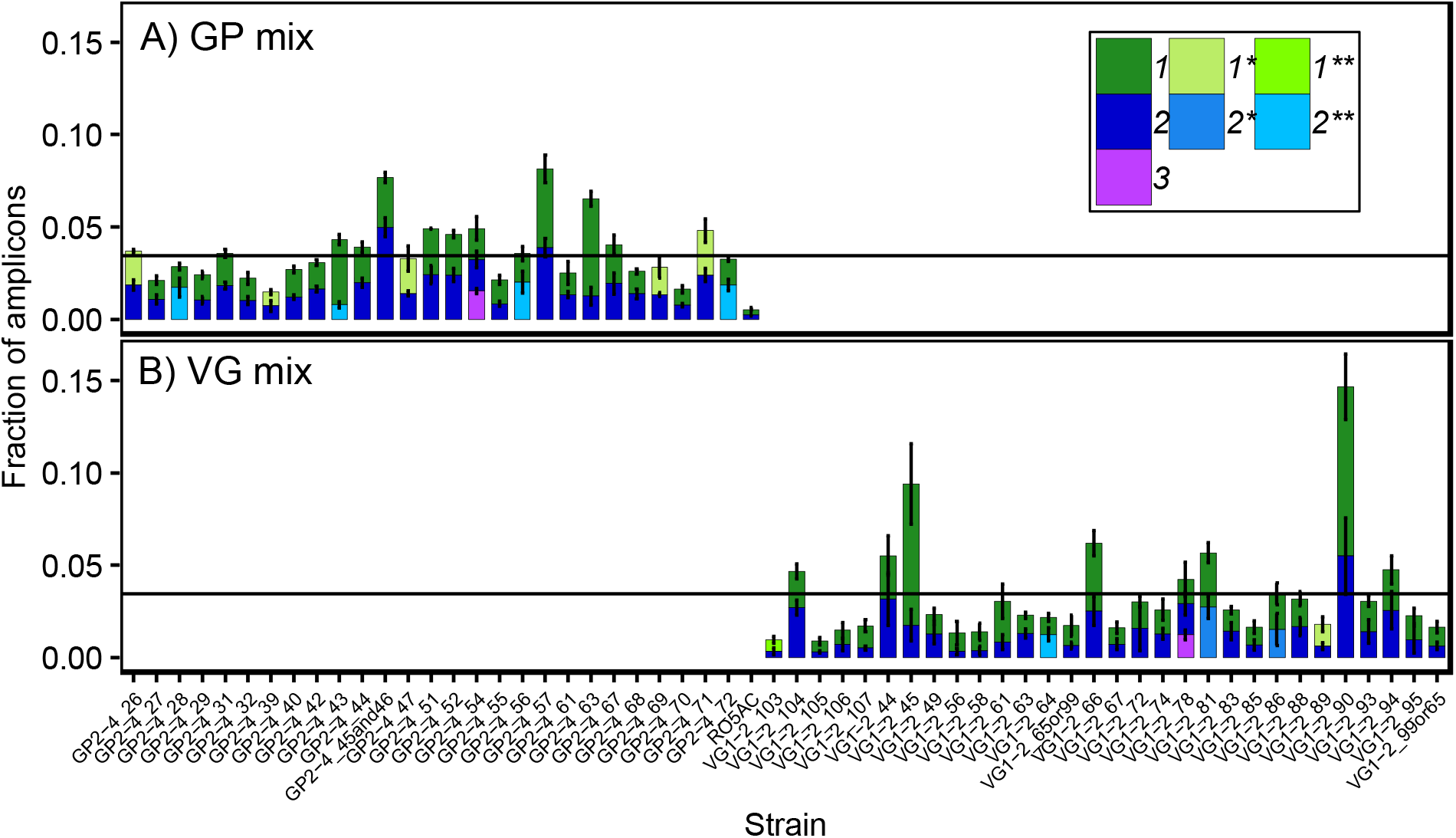
Metabarcoded relative abundance of strains alleles using the barcode loci *Sm_C12W1*. Shown are two mixed populations of *S. marinoi*. A) a mixture of 28 strains from Gropviken (GP), and B) 30 strains from Gåsfjärden (VG). Legend numbers indicate whether this is the first, second, or third allele in each strain, with colours/asterisks indicating if alleles are homologous in two (*) or three (**) other strains. For asterisk-marked alleles, the abundance has been partitioned between strains using differential equations. The horizontal line indicates the expected fraction of amplicons (allele 1+2) per strain, assuming even cell densities, Error bars show standard deviation, with N=3 technical PCR replicates for panel A, and N=4 for B. Data has been denoised using the settings for ‘Relaxed DADA2 on amplicons’ as outlined in the Supplementary Information.

Once the inaccuracies in the reference alleles were corrected using the confirmed sequences, merged amplicons that still did not match the expected allele sequences were investigated to determine the sources of error. This artifact search was limited to *S. marinoi* genotype samples with coverage >100 amplicons per locus. A detailed description of the analysis and its findings can be found in the Supplemental Information (Fig. S2, S3 and text). Briefly, 61-89% of amplicons contained sequencing-errors, with 39-76% failing to merge. 11 to 40% of merged amplicons mapped perfectly to known alleles, while remaining sequences were explained by erroneous base-calling (10-40%), off-target amplification (0.1-12%), and PCR chimeras (1-3%).

Most amplicon sequence-errors could be either corrected or filtered out using the denoising algorithm of DADA2 (Callahan et al. 2016). We finetuned the settings for the 523 bp *Sm_C12W1* locus to minimize false negative (alleles not predicted as real Amplicon Sequence Variants [ASV]) and false positive (any ASVs not corresponding to a biological allele) observations (Table S6). From mixed DNA samples, the most stringent settings evaluated failed to predict three out of 110 alleles, while 16 artefact sequences were retained as ASV, although they comprised only 0.03% of all merged sequences. By analysing the patterns of the remaining false positive ASVs across these samples, searching for linked alleles, we were able to iteratively identify the heterozygous alleles of three strains for which we lacked genotype sequencing data (GP2-4_32, VG1-2_65, VG1-2_99). The remaining false positives were annotated as either chimeras that DADA2’s *de novo* chimera filter failed to remove, or one bp sequencing errors from an abundant allele. Relaxing filtering settings, resulted in higher false positive ASVs (346, totalling 1.2% of all amplicons), but detection of all known alleles as ASVs (Table S6). Using the later settings on a mixed DNA sample, the *Sm_C12W1* alleles could be used to enumerate the abundance of 58 strains accurately (Fig. 4).

### Implementation of an intraspecific metabarcoding method in a microbial evolution experiment

The quantitative performance of *Sm_C12W1* as a metabarcoding locus was further evaluated in an artificial evolution experiment using *S. marinoi*. In the artificial evolution experiment, 30 strains from Gåsfjärden (VG) and 28 strains from Gropviken (GP) were density normalised (relative cell abundance of each strain was 3.4±0.89% based on cell counts using microscopy), pooled separately for each population, and subjected to 42 days (50-100 generations) selection with and without toxic copper stress. This data was used to evaluate the quantitative performance of *Sm_C12W1*’s ability to track strain abundances.

Despite our expectations of diploidy in the *S. marinoi* strains, there were signs of polyploidy in the experimental data. Some strains exhibited a skewed allele ratio closer to 1:3 (GP2-4_43, GP2-4_55, GP2-4_63, VG1-2_45, and VG1-2_61; Fig. 5) suggesting that they could be polyploids or that the PCR reaction amplified alleles with slightly different efficiencies. The combined allele abundance of strains was sometimes over-represented two- to three-fold (e.g., GP2-4_57, VG1-2_45, VG1-2_90; Fig. 4) while others were underrepresented at a similar magnitude (VG1-2_103, VG1-2_105, VG1-2_56; Fig. 4). This could not be explained by differences in cell density (R^2^=0.0003, slope=-0.0059, N=58), again suggesting ploidy differences, or different DNA extraction efficiencies between cells of strains. Despite these subtle deviations from expectations, the barcode *Sm_C12W1* could be used to identify all strains at close to the expected relative abundances, most often with strains two alleles at close to a 1:1 ratio (Fig. 4). As the populations evolved through selection on strain diversity, one or two strains ultimately became dominant in each treatment, and during this process, their allelic ratio was conserved at 1:1 (Fig. 5). These observations show that relative changes in allele frequencies should reflect changes in the relative abundance of strains.

**Figure 5:**
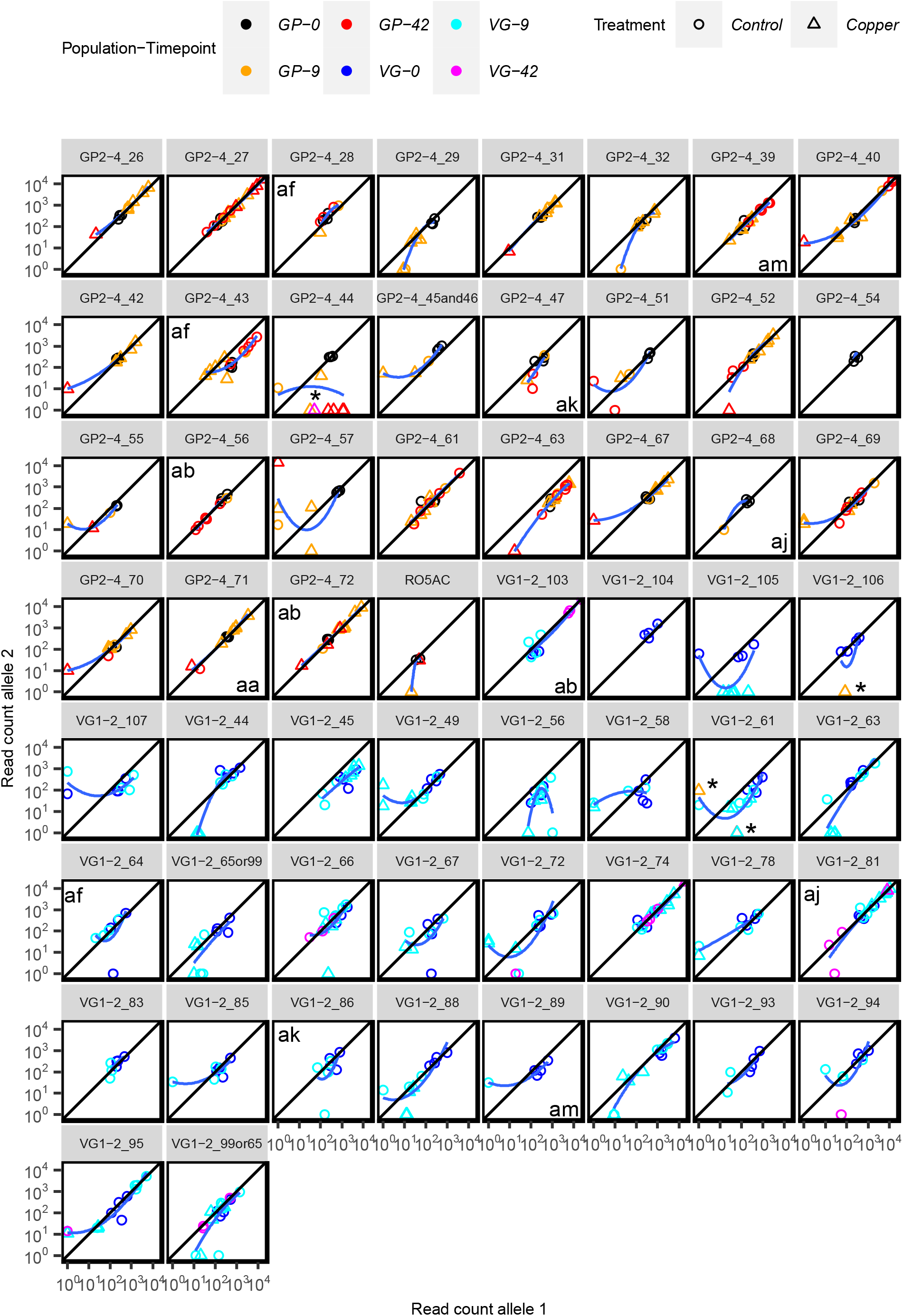
Correlations between heterozygous allele of 58 strains of *S. marinoi* across the 42-day selection experiment. For visibility, data has been transformed (x +1) with removal of 0+0 cases. Shared homologous alleles are indicated by alphabetical indexes in the corner of each allele (i.e., lower right equals allele one, upper left allele two), and their abundances have been allocated between strains using differential equations. All such equations assume a 1:1 ratio between alleles, except allele two in GP2-4_43, where a ratio of 0.15 is used based on the genotype sample. The black line across panels corresponds to the expected 1:1 ratio between heterozygous alleles. The blue line is a second order polynomial function fitted to the data, with shaded area corresponding to 95% confidence interval. Data has been denoised using the settings for ‘Relaxed DADA2 on amplicons’ as outlined in the Supplemental Information. Asterisks highlight false-positive observations that the denoising pipeline failed to remove. Fig. S4 shows the same figure without denoising (i.e. ‘exact matches’ settings in Table S6).

Throughout the experiment, most strains experienced negative selection. At around 10 ASV observations per sample, DADA2’s algorithm started filtering out observations (compare Fig. 5, with non-denoised sequence matches in Fig. S4), indicating that false-negative artefacts were created. However, DADA2 effectively removed all but four false-positive observations in the experiment (allele of GP population seen in the VG population, and vice versa: Tables S6). The remaining four false positives appeared to be chimeric, as did allele GP2-4_44#1, which displayed a skewed allelic ratio representing up to 5% (>1,000 copies) of amplicons in several replicate samples, while GP2-4_44#2 was not detected at all (Fig. 5). As the first 316 bp of this allele are identical with GP2-4_27#1, and the last 354 bp with GP2-4_27#2 (the two alleles of the most dominant strain in the same saples), there is a central 147 bp region where a PCR chimera from these two alleles would produce GP2-4_44#1. In addition to enabling chimera detection in situations like the one above, heterozygosity provided an additional analytical advantage, enabling differential equations to accurately partition alleles that were shared between two or three strains (Fig. 4 and 5). These results show that heterozygosity can be used to both filter out chimeric observations, and to enable improved quantification of strains, even if they lack a unique allelic marker.

## Discussion

By scanning the entire genome for suitable loci, we illustrate how the Bamboozle approach can identify hyper-variable barcodes with intraspecific resolution. In the case of both *S. marinoi* and *C. reinhardtii*, most of the genome (>90%) was not suitable for barcoding and was effectively filtered out by the algorithm. Of the remaining loci fulfilling the criteria for barcodes (Kress and Erickson 2008), only a fraction contained sufficient allelic richness to provide allele-rich markers for strain quantification, highlighting the power of a whole-genome scan approach for development of novel intraspecific barcodes. Using a combination of original (54) and published [17: (Flowers et al. 2015; Ness et al. 2016)] WGS datasets, we illustrate how such intraspecific barcodes can be designed from either publicly available sequencing data, or through the generation of new data for a species or population of interest.

Within our amplicon sequencing data, many amplicons (∼40%) did not correspond to any known allele in our database. Therefore, we attempted to identify the sources of these artefacts, and how to address them. This was done in order to make recommendations for future studies using workflows similar to ours.

First, we note that the 523 bp *Sm_C12W1* locus, paired with Illumina MiSeq’s 2×300 bp sequencing technology, creates some challenges that may not be encountered in shorter loci. For example, we could not use DADA2 as a stand-alone tool to process reads since it truncated 100% of all amplicons during merging (Table S6). BBMerge performed better but still failed to merge 78% of all reads, due to the short overlapping region and low base-call quality at the 3’ end of reads. By pairing BBMerge with DADA2, we found a flexible approach that could correct 99-99.97% of all sequencing errors in reads that had merged, with only spurious false-positive allelic observations (Fig. 5). Although a shorter locus is preferable and will likely perform better than this, our analysis shows that the tools available to make robust biological inferences from noisy amplicon data extend to 500 bp barcode loci, albeit with a loss of four-fifths of the sequence data. If future studies cannot identify window sizes smaller than 500 bp, accurate long-read sequencing of barcodes (e.g., PacBio’s Circular Consensus Sequences technique (Rhoads and Au 2015)), or paired end 300 bp sequences from the newer platform Illumina NovaSeq 6000 could be options worth considering. Long-read sequencing, in combination with DADA2’s error correction, has enabled differentiation between pathogenic and benign *E. coli* strains using single SNP markers and metabarcoding of the whole ∼1450 bp 16S rRNA gene (Callahan et al. 2019). Such a sequencing approach opens up for loci of several kb to be used as intra-specific barcodes.

In contrast to sequencing errors, PCR chimeras provided a more problematic type of artefact. We found that PCR chimeras could not be removed effectively using software built for processing metabarcoding data, such as Mothur (Rognes et al. 2016; Schloss et al. 2009), UCHIME2 (Edgar 2016), or DADA2s chimera removal algorithm (Callahan et al. 2016). Exacerbating this issue is the potential for ‘perfect fake’ sequences (chimeras identical to another, non-chimeric sequence in the dataset (Edgar 2016)). Out of the 110 alleles in *Sm_C12W1,* we identified several cases of such ‘perfect fakes’ (e.g., allele 1 in GP2-4_44; Fig. 5), and these instances should become progressively more numerous if more strains and alleles are included in the experiments and analysis. However, we could identify these PCR chimeras in our experimental design primarily by looking at deviation from the expected heterozygous allelic ratios of individual strain (Fig. 5). Depending on the research question and experimental design, chimeras can be more or less effectively filtered out using similar approaches in future studies. If chimeras risk causing erroneous biological interpretations from the data, approaches to reduce the number of chimeras produced includes optimising a low PCR cycle threshold (Smyth et al. 2010) or using chimera-free PCR kits, such as emulsion PCR (Williams et al. 2006).

The allele phasing step (upstream of Bamboozle) also created some problems, as it could not handle cases of triploidy (Fig. 4). Haplotype phasing of polyploids is non-trivial and benefits greatly from long-read data (Abou Saada et al. 2021). However, as long as polyploidy is accurately identified (e.g., by amplicon sequencing of each strain’s barcode individually), it should not provide any issues with downstream analysis of strain abundances and may even aid in providing increased allelic diversity for quantification purposes. Consequently our evaluation of Bamboozle as both a tool to identify intra-specific barcodes suggest it can accurately locate such hypervariable regions. Our quantitative analyse show that with appropriate choice of barcode lengths, PCR kits, sequencing platforms and denoising strategies, amplicon sequencing of intra-specific barcodes can be used to track the abundances of large numbers of strains from mixed DNA samples.

## Conclusions

We built and employed the Bamboozle tool to develop metabarcoding loci that provide intraspecific resolution in the diploid diatom *S. marinoi*. This allowed us to identify and quantify the abundance of 58 environmental strains from mixed DNA sample. We illustrated the usefulness of such an intraspecific barcode locus in an artificial evolution experiment where we tracked the abundance of all strains during co-cultivation. Although further tests are needed to assay performance in new populations and environmental samples, as well as the amplicon sequencing performance of the *C. reinhardtii* loci, our preliminary tests suggest that intra-specific metabarcoding can simplify or enable novel studies of microbial ecology and evolution in natural and artificial settings. This is not limited to artificial selection experiments like ours but possibly also to determine genotype diversity in natural populations, studies of ancient DNA from sediment cores (Adams et al. 2019; Härnström et al. 2011), mesocosm incubations of natural phytoplankton communities (Scheinin et al. 2015; Tatters et al. 2013), effects of predation (Sjöqvist et al. 2014), nutrient competition within (Collins 2011) and between species (Descamps-Julien and Gonzalez 2005), or other drivers that are challenging or impossible to observe in mono-culture experiments (Baert et al. 2016; Collins and Schaum 2021). Using the haploid chlorophyte *C. reinhardtii*, we illustrate how the Bamboozle tool should, if possible, be able to identify loci in both haploid and diploid organisms with a reference genome and whole genome sequencing data from multiple genotypes. Our comparative analysis of barcoding loci identified in *S. marinoi* and *C. reinhardtii* suggests that much of the genome is unsuitable for barcode design, and that different species have different optimal barcode loci. Consequently, Bamboozle’s strategy of scanning the entire genome of a species is a suitable approach to identify novel barcodes for evolutionary studies.

## Supporting information

All supplemental information and figures

## Acknowledgments

The authors would like to thank André Soares and Vilma Canfjorden who have developed additional functionality of the Bamboozle tool. Olga Kourtchenko, Lara Hoepfner, and Nora Klasen, assisted in experiments or DNA sample preparation. The manuscript benefited from revision suggestions and thoughtful comments by Kerstin Johannesson.

## Data availability

The code for Bamboozle is available at https://github.com/topel-research-group/Bamboozle and the WGS and amplicon sequencing data has been deposited at NCBI under BioProject PRJNA939970.

## Materials and Methods

### Strain retrieval and whole genome sequencing

Individual strains of *S. marinoi* were germinated and isolated from sediment resting stages using standard micropipetting techniques (Härnström et al. 2011). Surface sediment was collected from two semi-enclosed inlets of the brackish Baltic Sea, where one (Gåsfjärden [VG]; 57°34.35’N 16°34.98’E) has been exposed to historical copper mining (Söderhielm and Sundblad 1996) while the other (Gropviken [GP]; 58°19.92’N 16°42.35’E) has not. A total of 69 and 55 strains were isolated from VG and GP, with 88% and 94% survival, respectively. Strains were cultured in locally-sourced seawater (salinity 7), which had been sterile-filtered (Sarstedt’s [Helsingborg, Sweden] 0.2 mm polyethersulfone membrane filter), and amended with f/2 nutrients (Guillard 1975) and 106 µM SiO_2_. Cultures were continuously screened for contamination by other microalgae, and auxospore formation or bimodal cell sizes, which indicate sexual inbreeding in *S. marinoi* (Ferrante et al. 2019), and such cultures were discarded.

DNA extraction was performed within one month of revival from resting stages in a total of 28 strains from GP and 30 from VG. Cultures were made temporarily axenic via a combination of mechanical cleaning (triple washing in 20 µg mL^-1^ Triton X-100 media and cell collection on a 3.0-µm polycarbonate filter), followed by a five-day antibiotic cocktail treatment (90 µg mL^-1^ Paromomycin, ten µg mL^-1^ Ciprofloxacin, and 40 µg mL^-1^ Cefotaxime). On day six, cells were collected via centrifugation in 50 mL Falcon tubes (10 000 x g, for ten min.), flash-frozen in liquid nitrogen, and stored at −80°C. DNA was extracted using a CTAB-phenol–chloroform protocol as described in (Godhe et al. 2001), but with additional RNA digestion during cell lysis (65°C for 60 min. using 1 mg RNaseA mL^-1^ CTAB buffer). DNA yield was quantified using Qubit (Thermo Fisher Scientific). In two subsequential attempts, strains VG1-2_65 and VG1-2_99 did not survive the antibiotic treatment, and extractions yielded insufficient amounts of DNA for PCRs.

Whole genome sequencing was performed on the remaining 56 *S. marinoi* strains using ½ lane of an Illumina NovaSeq S4 flowcell. Library preparation was done using Nextera’s PCR-based protocol with enzymatic fragmentation. This resulted in 722.82 Mreads (2×150 bp paired-end).

### Data preparation for the pipeline – S. marinoi

Trimming of reads was performed on the whole genome sequencing data to remove the first 15 bases of each read. The trimmed reads were mapped to the *S. marinoi* reference genome version 1.1.2 (https://albiorix.bioenv.gu.se/Skeletonema_marinoi.html) using Bowtie2 version 2.3.4.3 (Langmead and Salzberg 2012), and the resultant SAM files were converted to sorted BAM files and indexed using SAMtools version 1.9 (Danecek et al. 2021). Variant calling was performed using GATK version 4.1.8.0 (Van der Auwera and O’Connor 2020), following the Best Practices recommendations on the tool’s website, and the resultant VCF files were indexed using BCFtools version 1.10.2 (Danecek et al. 2021). In addition, SAMtools version 1.10 was used to phase the BAM files, and GATK was re-run on each of these phased BAM files individually. These phased files were indexed as described above.

### Data preparation for the pipeline – C. reinhardtii

Sequencing data from the strains of interest was downloaded from either the NCBI Sequence Read Archive (SRA; downloaded using the `prefetch` and `fastq-dump` tools from the SRA toolkit version 2.9.6 (Leinonen et al. 2010)) or the European Nucleotide Archive (ENA; downloaded using the `enaDataGet` tool from enaBrowserTools version 1.6 [https://github.com/enasequence/enaBrowserTools]) (Table S4; retrieved 22 February 2021). PCR duplicates were removed using the filterPCRdupl Perl script version 2.3 (https://github.com/linneas/condetri), then trimmed with Cutadapt version 3.2 (Martin 2011), using a quality threshold of 30 and a post-trimming length threshold of either 90 (ENA reads) or 45 (SRA reads).

The trimmed reads were mapped to a reference consisting of the following sequences from *C. reinhardtii*: the contig-level assembly of the reference nuclear genome version 5.5 (accession no. ABCN00000000.2); the mitochondrial genome (accession no. NC_001638.1); the plastid genome (accession no. NC_005353.1); and the mating type minus locus (accession no. GU814015.1). Mapping was performed using Bowtie2 version 2.3.4.3 (Langmead and Salzberg 2012), with the resultant SAM files being converted to sorted BAM files and indexed using SAMtools version 1.10 (Danecek et al. 2021). Variant calling and VCF indexing were performed as described above for *S. marinoi*.

### Bamboozle workflow

Barcodes for *S. marinoi* were obtained using commit 75467da of the Bamboozle main branch (except for *Sm_C2W24*, which was obtained using commit ae28b1b). Barcodes for *C. reinhardtii* were obtained using commit db64632 of the `matt_improvement` branch. The latter commit has the fourth and fifth steps of the pipeline reversed (i.e., windows with irregular coverage are excluded prior to window merging), along with other minor improvements.

The input files (reference genome FASTA files, plus BAM and VCF files [and respective indexes] for each strain) were prepared as described above. Bamboozle was run in both species with a `–window_size` parameter of both 500 and 300, and a `–primer_size` parameter of 21.

### Development of the bioinformatic pipeline

The predicted barcodes were first visualised against the reference genomes using IGV version 2.6.3 (Robinson et al. 2011) to look for anomalies. In an early iteration of the pipeline (commit ae28b1b) without coverage depth filters (steps 1 and 5, Fig. 1), many loci contained obvious errors in the conserved primer sites, or spanned regions of low read coverage (e.g., *Sm_C2W24*) suggesting the pipeline could not effectively process regions with large indels (e.g. micro- or mini-satellites). This issue has largely been resolved by adding the coverage depth filter, as evidenced by the reduction of identified barcodes in *S. marinoi* from 245 to 4.

Additional adjustments of the pipeline used for analysing both *S. marinoi* and *C. reinhardtii*, included extending the code to allow analysis of haploid genomes, and steps 4 and 5 of the pipeline were switched (i.e., windows with irregular coverage are excluded prior to window merging), in order to avoid excluding potentially informative windows without coverage issues.

### In silico annotation of intraspecific barcodes

The gene models of the predicted barcodes were retrieved from version 5.5 of the *C. reinhardtii* reference genome (https://ensembl.gramene.org/Chlamydomonas_reinhardti) and version 1.1.2 of the *S. marinoi* reference genome (https://albiorix.bioenv.gu.se/Skeletonema_marinoi.html). Functional domains were annotated using NCBI’s CD-Search webtool (Marchler-Bauer and Bryant 2004). When gene models lacked a functional annotation, putative functions were assigned by a homology search using BLASTp against the NCBI database (Altschul et al. 1990), and comparison of functional domain structure.

To assess primer specificity outside our Baltic Sea strains of *S. marinoi,* we mapped transcriptional data from three strains from global culture collections (Keeling et al. 2014), and manually looked for variants within the predicted conserved regions. A targeted BLASTn search was used to identify orthologous genes in three species within the order *Thalassiosirales* with reference genomes, namely: *Thalassiosira oceanica* (Lommer et al. 2012), *Thalassiosira pseudonana* (Armbrust et al. 2004), and the closely related *Skeletonema subsalsum* (Sarno et al. 2005) using an in-house draft genome for the latter (Pinder et al. unpublished data).

### Confirmation of the bioinformatic predictions

Illumina amplicon sequencing was used to assess the sequence accuracy of each of Bamboozle’s locus predictions in the diploid *S. marinoi,* and Sanger sequencing for the haploid *C. reinhardtii*. Primers (standard de-salted oligos) were designed for the 21 bp conserved regions flanking the variable region (Table S5) and extended with Illumina adapter sequences for *S. marinoi*. *rbcL* was used as a positive control of amplification in all diatoms (Guo et al. 2015), and 16s or 18S rRNA across other phytoplankton taxa (Stoeck et al. 2010; Sundberg et al. 2013). The barcodes were amplified from 100 ng DNA and a final primer concentration of 0.1 µM using the Phusion® High-Fidelity PCR Kit (Thermo Scientific) with individually optimised annealing temperatures between 62 and 65°C. A one-step PCR reaction, with adapter extended primers, was run for 30 cycles, with 5 s denaturation (98°C), 5 s annealing, 30 s extension (72°C), and a final 5 min extension. The evenness of amplification across strains was qualitatively assessed using gel electrophoresis. Strains with poor barcode amplification were re-run with either increased DNA concentration or up to 35 PCR cycles. Species specificity of the primers was assessed using DNA from two strains of *Skeletonema subsalsum*, as well as a mixture of other common phytoplankton species from the Baltic Sea (see Supplemental information). The *C. reinhardtii* primers were only tested on two other green algae strains; *Chlamydomonas* sp and *Microglena* sp.

For the 56 *S. marinoi* strains, the barcode alleles were sequenced individually using 2×301 bp reads on 1/10 of a lane on the Illumina MiSeq platform with v3 chemistry (expected 1.8 million reads in total). The four PCR reactions were run independently for each barcode, and the products pooled based on volume. Further library preparation, including amplicon size selection >300 bp, Nextera dual-indexing, and amplicon sequencing were performed at SciLifeLab (NGI Stockholm), according to the manufacturer’s instructions.

### Identifying sources of error in the amplicon sequencing results

Paired end reads were first quality filtered and trimmed using Cutadapt version 3.2 to remove adapter and primer sequences, with a quality threshold of 28 and a minimum read length of 180 bp. The resultant trimmed reads were merged using BBMerge version 38.86 (Bushnell et al. 2017). Amplicon sequences from the individually strains that did not match the two expected allele sequences were investigated to determine the sources of error. Chimera detection was initially performed using the `chimera.vsearch` function of Mothur version 1.47.0 (Schloss et al. 2009). However, as this highlighted fewer chimeras than we knew to exist through manual inspection, we instead switched to the `uchime2_ref` function of Usearch version 11.0.667 (Edgar 2016). This was run with the options ‘`-strand plus -mode sensitive’` against a database containing all alleles expected to appear in that strain. Identification of truncated sequences, sequences containing Ns, off-targets, and localisation of sequencing errors and SNPs along the amplicons, was performed using a custom Python script, incorporating Bowtie2 version 2.3.4.3 (Langmead and Salzberg 2012), SAMtools version 1.12, and BCFtools version 1.12 (Danecek et al. 2021).

### Performance of intraspecific metabarcoding in a selection experiment

The quantitative performance of *Sm_C12W1* barcodes locus was validated in a selection experiment using the 58 *S. marinoi* strains. A detailed description of this experimental design and the results will be published elsewhere (Andersson et al, unpublished). Briefly, the single strain cultures (non-axenic) were pooled back to their original populations (Gropviken: GP and Gåsfjärden: VG) at even densities, and maintained in exponential growth phase via a semi-continuous dilution scheme as outlined in Andersson et al. (2020) through dilutions every third day for a total of 42 days. Five 100 mL replicates per population were grown with toxic copper stress of 8.65 µM CuSO_4_, and five replicates were grown without toxic stress. The dilution bottleneck was ∼10,000 chains of cells per replicate. Cultures were harvested at the start of the experiment, after nine, and 42 days of selection. DNA was extracted from experimental samples as for the WGS preparation outlined above, but without the axenic treatment. Barcodes were PCR amplified from each replicate timepoint. For the single time 0 sample, two technical DNA extractions replicates × two PCR replicates were processed and analysed in parallel (i.e., N=4 technical replicates). The selection experiment samples were indexed and pooled together with the individual genotypes, on the same MiSeq run using 9/10 of the read capacity (i.e., an expected 16.2 million reads).

The individual strains’ genotype samples were used to create a database of all strains’ alleles, and amplicons from the selection experiment samples were compared against this database (script available at: https://github.com/topel-research-group/Live2Tell). As the experimental samples contained mixtures of strains at variable densities, and allele calling was expected to require single SNP resolution across long amplicons (up to 523 bp), we explored avenues to denoise the data. Initial attempts to use the nf-core/ampliseq pipeline version 2.4.0 (Straub et al. 2020) failed to detect all expected alleles. Running DADA2 version 1.16.0 (Callahan et al. 2016) – a component of the nf-core/ampliseq pipeline – as a standalone program for identifying amplicon sequence variants (ASVs) proved more successful. Different settings and combinations of steps were iteratively explored (described in detail in Supplemental information). We used four criteria to evaluated the quality of the output including: 1) how many known allele sequences were <u>not</u> identified as ASVs (false negatives); 2) how many unexpected ASVs were outputted (false positives); 3) what fraction of total raw reads assembled into amplicons matching either known alleles or false positives; and 4) if false positive or false negative observations risked affecting the biological interpretation of the artificial evolution experiment.

The relative abundances of strains were quantified based on the number of amplicons matching its two alleles. In this step, a set of differential equations were used to parse out the observations of alleles shared between strains. In general, the proportion of shared allele *i*_1_ belonging to a specific strain x, y, and z was computed individually for each sample based on strains second unique allele (*x_2_, y_2_,* and *z_2_*):

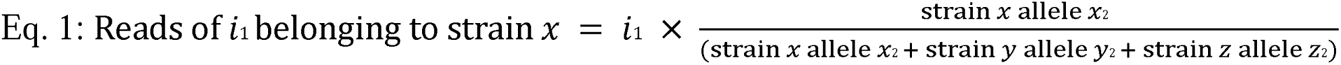

The two heterozygous allele counts were then summed up to represent the abundance of each strain, with the third allele of triploid strains omitted. Amplicon counts per strain were simultaneously normalized to relative abundance (RA) as:

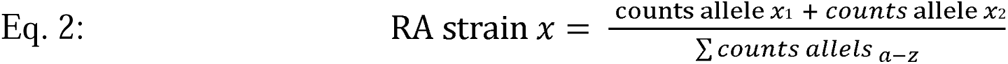

To validate the approach, we assessed how closely the barcode enumeration of strains matched with microscopic cell counts in the time zero samples and how closely the abundance of each strain’s two alleles correlated throughout the selection experiment.

