## Supplementary material for "Bamboozle: A bioinformatic tool for identification and quantification of intraspecific barcodes": All supplemental information and figures

#### Detailed in silico annotation of intra-specific barcode features

All four barcode-containing genes in *S. marinoi* (Fig. 3A) encoded proteins predicted to interact physically with other biomolecules. *Sm_C2W24* was inside Sm_t00004275-RA, a gene with a unique domain structure that we could not find any homologs in the NCBI database (analysis performed December 12th, 2021). It contains three repeats of a Small Ubiquitin-like Modifier (SUMO) domain, involved in post-translational protein modification, with the primer sequences anchored in the second and third repeat (Fig. 3A). SUMOs proteins are well characterized in humans (Verger et al. 2003), yeast (Hofmann et al. 2000), and Arabidopsis (Morrell and Sadanandom 2019). In these species, the SUMO domain is only presented in a single copy in a short gene, and the ca. 9-11 kDalton peptide is covalently attached to primarily lysine residues of protein to modify their function. Like ubiquitin, the SUMO modification can serve a multitude of different functions, including targeting proteins for degradation, or changing the activity of transcription factors. The centre of the barcode contains a nine bp repeat encoding Alanine- Asparagine- Glutamic acid [occasionally replaced by Glycine] which we refer to as the AN(E) repeat. The AN(E) repeat of Sm_t00004275-RA has no homologs amongst phytoplankton in NCBIs current database, but it has similarity to viral proteins in *Plasmodium spp, Vibrio casei,* and *Cotonvirus japonicus*, which all have genes with an 18mer repeat structure yielding peptides with similar residues (e.g., TNEANE, NETNEV, and ANDAGE).

*Sm_C12W1* was located inside a pentatricopeptide (RNA binding) repeat region of Sm_t00009768-RA. The closest BLASTp match in the NCBI database was a ‘maturation of rbcL mRNA’ protein in *A. thaliana* (AT4G34830.1). However, these genes are likely not orthologous, as the *S. marinoi* genome has more than 20 genes similar to Sm_t00009768-RA and AT4G34830.1*,* and out of these Sm_t00011335-RA has higher amino acid similarity to AT4G34830.1. Consequently, Sm_t00009768-RA may be involved in mRNA maturation or types of other RNA interactions, but perhaps not specifically for rbcL.

*Sm_C12W2* was located inside one of two YheB membrane domains of Sm_t00009769-RA*.* This gene is relatively conserved in *T. pseudonana* (Thaps3_5794; 72% coverage and 69% identity at the nucleotide level), but has three and four mismatches in the forward and reverse primer sites, respectively (Table S3). We also found a protein homologous to Sm_t00009769-RA in *T. oceanica* (THAOCEA1_01345), but nucleotide similarity was eroded to the point that we could not accurately identify the primer sites (Table S3).

*Sm_C16W4* was located inside a WD40 repeat of Sm_t00008465-RA (Fig. 3A), whose protein is similar to microtubule-associated ribosome biogenesis protein YTM1 in *S. cerevisiae*. Unfortunately, the *SM_C16W4* primer set (Table S2) did not amplify well once adapters had been added to primers, likely due to increased dimerization or a modified secondary structure (Fig. 3B).

#### Identification of sources of amplicon errors of barcodes in S. marinoi

Despite quality filtering (3′ Illumina adapters removed, 3′ quality filter of Phred score 28, minimum post-trim read length 200), sequencing errors were present in most of our data (affecting up to 75% of all amplicons). We tried to increase the Phred quality score filter from 28 to 30, but this resulted in 98% of all reads failing to merge, which was unacceptably high.

Secondly, despite checking the primer sequences using BLASTn against the *S. marinoi* reference genome prior to PCR amplification and sequencing, attempts to amplify some of our loci resulted in substantial amounts of off-target amplification and primer dimer formation. In our study, this seemed to particularly affect *Sm_C12W1* and *Sm_C16W4*; the *Sm_C16W4* off-target/primer dimer was short and removed during size selection or adapter filtering, but *Sm_C12W1* escaped these steps and merged into amplicons, accounting for an average 20.97% of merged amplicons (Fig. S2). The terminal regions of the 358 bp off-target sequences in *Sm_C12W1* don’t match the full length of the primer sequences, and BLASTn does not map the primers sequences to these regions. Yet the full off-target sequences have strong BLASTn matches at nine different contigs in *S. marinoi*, with themselfe have five to ten-fold higher coverage depth than the genomic average, suggesting they are repetitive element occurring throughout the genome. This likely explains why the region amplified despite having a relatively poor predicted annealing of the primers. As with sequencing errors, off-targets lower the number of amplicons per sample, but are relatively simple to remove bioinformatically. Consequently, off-targets are unwanted but of little concern to downstream analyses.

The third issue identified with Bamboozle was the appearance of unexpected triploid strains in our experiment. The default expectation of the phasing and variant calling software used for the *S. marinoi* study was diploidy, and this was the ploidy defined in Bamboozle. Furthermore, as the coverage filter employed looks at the coverage across the genome on a per-contig level, it cannot detect duplication of entire contigs/chromosomes corresponding to aneuploidy, or the presence of full polyploidy. Upon subsequent manual inspection of the WGS results, we identified some of our strains as triploid.

The fourth and most serious problem was the presence of PCR chimeras that the bioinformatic chimera filters failed to remove. The chimera removal software we initially employed (the chimera.vsearch command of Mothur version 1.47.0) only identified an average of up to 0.3% of all merged reads in each barcode as being chimeric. However, as chimera removal software often fails to detect chimeras amongst highly similar sequences (>95%), which included almost all of our alleles, we manually examined some of our results and determined that chimeras were indeed being missed by the software. We thus tried a different tool (the uchime2_ref command of Usearch version 11.0.667_i86linux32), whose 'sensitive' mode produced a result more in line with our manual observations. It should be noted, however, that this mode of the software can produce false positive results. This analysis gave chimera values of between 0.77% and 4.02% of merged reads (Fig. S2), but manual annotation found that chimeras were still present in the filtered data. As chimeras will duplicate during each PCR cycle, they can rapidly generate many exact copies of an artefact amplicon, in contrast with sequencing errors that are, although not completely randomly created, at least more evenly distributed (Fig. S3). Chimeras will also only affect SNP positions across the read. At the same time, sequencing errors are expected to affect SNPs and non-informative sites equally, and the chimeric PCR mechanism is indistinguishable from genomic recombination that generates natural diversity within populations. Consequently, chimeras were strongly overrepresented in generation of false positive observations in some strains (Fig. 5 and S5). However, we found that the diploid nature of vegetative *S. marinoi* strains provided opportunities to detect these chimeras, as well as other artefacts (Fig. 6), and they can be masked in various ways when amplicon counts are translated into cell or strain counts.

*Attempted optimization of DADA2 parameters*

As our amplicon sequencing data contained various errors (see section *Identification of sources of amplicon errors of barcodes in S. marinoi* above), we attempted to denoise the data using DADA2. This pipeline was applied to the 43 amplicon sequencing samples of the *Sm_C12W1* barcode locus on mixtures of strains with various strain complexity (same data as shown in Fig. 4 and 5 of main manuscript). The results of this analysis are presented in Table S6.

All samples were first quality controlled using Cutadapt, as described in the Materials and Methods section of the main text, cutting the barcode length from 523bp to 484bp with primer sequences removed. To establish a baseline against which to compare DADA2’s performance, we merged the paired-end reads using BBMerge (default parameters), with no denoising (**Exact matches**). While this returned all 110 *Sm_C12W1* expected alleles, it also returned 149,753 false positives (defined here as those ASVs with a length +/- 12bp the expected 484bp barcode length, and containing the 5’ and 3’ regions identical among the expected alleles, thus filtering out truncated amplicons, off-targets, and non-*Sm_C12W1* amplicons).

We next attempted to run the reads through the nf-core/ampliseq pipeline, which includes a DADA2 step for denoising and read merging (**ampliseq**). However, this returned none of the *Sm_C12W1* alleles. Upon closer inspection, this was due to the minOverlap and maxMismatch parameters of DADA2’s mergePairs function being set to the default values (12 and 0, respectively). These parameters are not adjustable within the nf-core/ampliseq pipeline. Thus, we attempted to run DADA2 as a standalone program, keeping many of the parameter values used by nf-core/ampliseq, but adjusting the mergePairs parameter values to match the BBMerge defaults (minOverlap 0 and maxMismatch 20), as well as setting removeBimeraDenovo’s minFoldParentOverAbundance parameter to 90 (default 1.5) (**DADA2 with merging**). While this returned all but one of the expected alleles, we also obtained almost a thousand false positives, as well as experiencing unexpected drop-out of several expected alleles in some samples.

Despite attempts to solve these issues within DADA2, we ultimately opted to merge the reads in BBMerge, before passing the merged reads to DADA2 as ‘single-ended’ reads. In addition to running the previous iteration of the pipeline on these pre-merged reads (**Relaxed DADA2 on amplicons**), we also ran the pipeline with DADA2 default settings, but implementing an error inflation step to compensate for the adjustment of quality scores in the read overlaps as performed by BBMerge (this additional step is recommended by one of DADA2’s developers on the software’s GitHub page) (**Stringent DADA2 on amplicons**).

### Supplemental Tables and Figures

Table S1: Genetic variability amongst 55 *Skeletonema marinoi* strains in common metabarcoding loci previously suggested having some degree of intra-specific resolution in the diatom order *Thalassiosirales*. Abbreviations – ITS: internally transcribed spacer; rbcL: ribulose-1,5-bisphosphate carboxylase/oxygenase large subunit; COI: cytochrome *c-*oxidase subunit 1.

| Gene | Coordinates | Length (bp) | SNPs | Strains with same genotypes (max 55) |
| --- | --- | --- | --- | --- |
| 18S rRNA | Sm_000009F:1494606-1496397 | 1700 | 1 | 52 |
| 28S rRNA^2^ | Sm_000009F:1497116-1497900 | 785 | 0 | 55 |
| (18S+ITS1+5.8S+ITS2)^1^ | Sm_000009F:1496358-1497139 | 800 | 8 | 46 |
| rbcL | Sm_plastid:99162-100634 | 1473 | 0 | 55 |
| COI^3^ | Sm_mitochondrion:2060-3562 | 1503 | 2 | 53 |

^1^ Partial. Proposed as an intra-specific or population marker loci for *Thalassiosirales* by Guo et al. (2015)

^2^ Used to resolve abundance of species within the *Skeletonema* genus using metabarcoding (Canesi and Rynearson 2016)

^3^ Proposed to have some intra-specific variation in *Skeletonema* genus by Yamada et al. (2017)

Table S2: List of all 26 barcode windows identified by Bamboozle, with the genomic coordinates, the sequence of the conserved regions, and the putative function of the gene containing the locus.

| Species | ID | Genomic coordinates | Total length (bp) | 5' end conserved | 3' end conserved | Gene model (annotation) |
| --- | --- | --- | --- | --- | --- | --- |
| *S. marinoi* | *Sm_C12W1* | Sm_000012F:1009438..1009953 | 516 | CCTCAAACCCCATCGAATACAAACTCTCCATATGCA | GCTTTGCGGAATCTTTGCGTGTGCATCGTGCCCATG | Sm_t00009768-RA (TPR-like |
|  |  |  |  |  |  | superfamily protein) |
| *S. marinoi* | *Sm_C12W2* | Sm_000012F:1011059..1011599 | 541 | GAACTCCAAAAAACACCCCTCCAATACGCAACAATCCCCGCAGTAGCCGCCTTCCTAGGCC | TTGTTTCAAAAGGTGGGACGTGTGGAATTGGATTTTTTGGTGGAAAGTGGATTGGGGTTTG | Sm_t00009769-RA |
|  |  |  |  |  |  | (Uncharacterized membrane protein) |
| *S. marinoi* | *Sm_C12W3* | Sm_000012F:1011134..1011634 | 501 | GCCGGAGTCAAAATGTTATTC | TTGGTGCAGATGGTGGCGTGG | Sm_t00009769-RA |
|  |  |  |  |  |  | (Uncharacterized membrane protein) |
| *S. marinoi* | *Sm_C16W4* | Sm_000016F:654027..654528 | 502 | ATCAAGTGTGTGTCATCTTTGA | ACTCTCCTCACGGGAAGTTGGG | Sm_t00008465-RA (ribosome |
|  |  |  |  |  |  | biogenesis protein WDR12) |
| *C. reinhardtii* (mt-) | *Cr_Chr2W4* | chromosome_2:5974497-5974828 | 332 | GCGCCGGCCCTGACTCTATCTTTTGAAAGTTCACGAGATGTAGGCAAAGAGC | CGCTGGCGGACGCGCTCAGCCGGAGCCTGCGGGAGCGCTACCCCGAAAGCAA | CHLRE_02g112150v5 (phospholipase B) |
| *C. reinhardtii* (mt-) | *Cr_Chr9W12* | chromosome_9:3429678-3429981 | 304 | CACCACGTCACACTGCCCTGGTCC | ATCGCACAGGTGGGTGCCGACGCG | CHLRE_09g389134v5 (unknown function (DUF563)) |
| *C. reinhardtii* (mt-) | *Cr_Chr9W13* | chromosome_9:3429715-3430015 | 301 | ACCTGGTGTCCATCCTGCAGG | CAGCTTACAAGGCGGGCGTGC | CHLRE_09g389134v5 (unknown function (DUF563)) |
| *C. reinhardtii* (mt-) | *Cr_Chr10W15* | chromosome_10:3895695-3895996 | 302 | GCGGCAACCGGTGCCCAGCGCG | CGTGGCCAAAGCGGTGCCGGCG | CHLRE_10g447800v5 (60kDa SS-A/Ro ribonucleoprotein?) |
| *C. reinhardtii* (mt-) | *Cr_Chr13W19* | chromosome_13:4163058-4163376 | 319 | GTGCGGCCGGGCGCGGGTAAGTGCCTGGCTGAGGTCGAC | GCGTCATCGACGTGCACATCCACCTCAATGAGCCCGGTC | CHLRE_13g592050v5 (allantoinase) |
| *C. reinhardtii* (mt-) | *Cr_Chr16W20* | chromosome_16:2930253-2930565 | 313 | TGCCACGCCGGATGCCATACACAGCATGCGCCC | TGAAGCTGCTGCCGGACGCCAAGTGCGTGAGCG | CHLRE_16g664050v5 (GRAM domain? DUF4782?) |
| *C. reinhardtii* (mt-) | *Cr_Chr17W6* | chromosome_17:1200452-1200762 | 311 | TATATGGCGCAGGAGGTGACGGTGGTCCAGG | CAGGCCAACGCCAAGCGCGACCGTGATGTGG | CHLRE_17g704950v5 (unknown function) |
| *C. reinhardtii* (mt+) | *Cr_Chr1W1* | chromosome_1:5482222-5482550 | 329 | GACCGTGACTACGGCATCTTCAACAAGATCCACCACGACATCGGCACCC | CTGCACAGATTCCCCACTACAACCTTGAGGAGGCTACCGAGGCCGTCAA | CHLRE_01g038600v5 (chloroplast glycerolipid omega-3-fatty acid desaturase) |
| *C. reinhardtii* (mt+) | *Cr_Chr1W2* | chromosome_1:5482306-5482615 | 310 | AGGGCTGAATCTGGGTCGGGTTGGGAAATG | AGAGCCCCGGCCCCCTGCCCACCCACCTGG | CHLRE_01g038600v5 (chloroplast glycerolipid omega-3-fatty acid desaturase) |
| *C. reinhardtii* (mt+) | *Cr_Chr1W3* | chromosome_1:5735287-5735592 | 306 | GGTCGCTGCCACACTTCACCCCGAGG | TCCTGACTGCTGGACCAACCGCTCGC | CHLRE_01g040900v5 (unknown function) |
| *C. reinhardtii* (mt+) | *Cr_Chr3W5* | chromosome_3:3981328-3981632 | 305 | CCACCTACCCCTCCCGCTCCGTGTC | CACAGACGAGGAGCGGGAGGAGGAG | CHLRE_03g171850v5 (unknown function) |
| *C. reinhardtii* (mt+) | *Cr_Chr3W6* | chromosome_3:5068965-5069267 | 303 | ACATGAGCGTTGCAACATGTGTA | GCTATGCTGTCCGTCGGTTTTTG | CHLRE_03g181050v5 (unknown function) |
| *C. reinhardtii* (mt+) | *Cr_Chr3W7* | chromosome_3:5075944-5076245 | 302 | TGGTTCACGGGCGGCACCGTGA | GCAGCGTAGTTTTGATATGGGC | CHLRE_03g181100v5 (hypothetical protein) |
| *C. reinhardtii* (mt+) | *Cr_Chr3W8* | chromosome_3:5075977-5076278 | 302 | TCCTCCACATGGCCCTTAGCCT | GCACATTCCGGTTCACCAGGTC | CHLRE_03g181100v5 (hypothetical protein) |
| *C. reinhardtii* (mt+) | *Cr_Chr3W9* | chromosome_3:6202804-6203115 | 312 | CGGCCGCACCTTGCCCGCCACCGCGGCGCCGC | ATCCTGGCCAAGTGGCTGGCTTGCAAGGCTGC | CHLRE_03g192350v5 (Kelch motif?) |
| *C. reinhardtii* (mt+) | *Cr_Chr3W10* | chromosome_3:7273555-7273862 | 308 | GCACGCAACGCGGCTGTTGGTCTCGCTG | GAAACGCATATAAACCTCCGATCGAATG | CHLRE_03g207250v5 (putative glutamine synthetase) |
| *C. reinhardtii* (mt+) | *Cr_Chr6W11* | chromosome_6:7174112-7174428 | 317 | GAGGTCAGGGTAGCTGCTGTGGCCTCGGACTTGAGGG | CCTTGGCGGTCAGCGTCATGTGCCCGGTTGGAAGCCC | CHLRE_06g297800v5 (hypothetical protein) |
| *C. reinhardtii* (mt+) | *Cr_Chr11W16* | chromosome_11:59098-59403 | 306 | AGATGTTGATGTTGATGGCGCCGAAC | TGTTGATGGACCTCAGCGGCCGCAGC | CHLRE_11g467528v5 (calcium channel) |
| *C. reinhardtii* (mt+) | *Cr_Chr11W17* | chromosome_11:59130-59458 | 329 | AAGTAGAACACGGCCAGGATGACCACGTCCAGCAGCAGCGGAATGGAGT | ACGCAGCGGATGAGTGTGTAGTTGCCGCGGCCCGACAGATCCAGGTAGC | CHLRE_11g467528v5 (calcium channel) |
| *C. reinhardtii* (mt+) | *Cr_Chr11W18* | chromosome_11:59181-59493 | 313 | AGCATGGTGTCCACCAGTTGCTGTGTGGGGTAA | AGGCCCACAACCACAAAGTCTATGATGTTCCAC | CHLRE_11g467528v5 (calcium channel) |
| *C. reinhardtii* (mt+) | *Cr_Chr17W21* | chromosome_17:1192804-1193104 | 301 | TGCCCTGGATCTTGTCGCGGC | AAGTGCACTGCACCGTGTGTG | CHLRE_17g704850v5 (adenine phosphoribosyltransferase) |
| *C. reinhardtii* (mt+) | *Cr_Chr17W22* | chromosome_17:1192828-1193181 | 354 | TGAGGAAGGGCAGCTCGATGACGCACGCCGCCTCCACCACCACACCGCCGGCCTTCTCTGTGGGGCGGTGAGGG | GGCGCATGTGAGACTTACTGACGAGGTTGATGCCGGCGGCAAGGGTGCCGCCGGTGGCAATCAGGTCATCAACC | CHLRE_17g704850v5 (adenine phosphoribosyltransferase) |

Table S3: Primer mismatches in other populations of *S. marinoi* and species within the *Thalassiosirales* order. Sequences were obtained from either the Marine Microbial Eukaryote Transcriptome Sequencing Project (*S. marinoi*; (Keeling et al. 2014), NCBI (*Thalassiosira pseudonana* (Armbrust et al. 2004), and *Thalassiosira oceanica* (Lommer et al. 2012)), or an in-house draft genome assembly (*Skeletonema subsalsum*). NA indicates that the genomic region of the barcode could not be identified using BLASTn, and ND indicates insufficient sequencing depth of transcriptomic data (<1 coverage across the primer site) to determine if mismatches were present. Gene model homologs in *Thalassiosira* included – *Sm_ C2W24*: NA; *Sm_C12W1*: THAPS3_24049; *Sm_C12W2*: THAPS3_5794, THAOCEA1_01345; *Sm_C16W4*:  THAPS3_22985, THAOCEA1_70515.

| Primer site | Adriatic Sea *S. marinoi* (strain FE7) | Adriatic Sea *S. marinoi* (strain FE60) | Narragansett Bay *S. marinoi* (strain skelA) | *S. subsalsum* (strain LO-03-75) | *T. pseudonana* (strain CCMP1335) | *T. oceanica* (strain CCMP1005) |
| --- | --- | --- | --- | --- | --- | --- |
| *Sm_C2W24-F* | 0 | 0 | 0 | NA | NA | NA |
| *Sm_C2W24-R* | 0 | 0 | 0 | NA | NA | NA |
| *Sm_C12W1-F* | 0 | 0 | 0 | 4 | 3 | NA |
| *Sm_C12W1-R* | 0 | 0 | 0 | 0 | 7 | NA |
| *Sm_C12W2-F* | 0 | ND | 0 | 1 | 3 | >10 |
| *Sm_C12W2-R* | 0 | 0 | 0 | 0 | 4 | >10 |
| *Sm_C16W4-F* | ND | ND | 0 | 5 | 7 | 7 |
| *Sm_C16W4-R* | ND | ND | 0 | 3 | 6 | 7 |

Table S4: Strains of *Chlamydomonas reinhardtii* used in this paper, their accession numbers, the database from which they were retrieved (either NCBI’s Short Read Archive [SRA], or the European Nucleotide Archive [ENA]), and the relevant citation.

| **Strain** | **Accession no.** | **Database** | **Citation** |
| --- | --- | --- | --- |
| **Mating type +** | | | |
| CC-2343 | SRX823807 | SRA | (Flowers et al. 2015) |
| CC-2344 | SRX823856 | SRA | (Flowers et al. 2015) |
| CC-2936 | SRX823810 | SRA | (Flowers et al. 2015) |
| CC-2937 | SRX823858 | SRA | (Flowers et al. 2015) |
| CC-3065 | SAMEA4731376 | ENA | (Ness et al. 2016) |
| CC-3071 | SAMEA4731378 | ENA | (Ness et al. 2016 |
| CC-3076 | SAMEA4731381 | ENA | (Ness et al. 2016 |
| CC-3086 | SAMEA4731384 | ENA | (Ness et al. 2016 |
| **Mating type -** | | | |
| CC-2290 | SRX823806 | SRA | (Flowers et al. 2015) |
| CC-2342 | SRX823805 | SRA | (Flowers et al. 2015) |
| CC-2931 | SRX823857 | SRA | (Flowers et al. 2015) |
| CC-2935 | SRX823809 | SRA | (Flowers et al. 2015) |
| CC-2938 | SRX823808 | SRA | (Flowers et al. 2015) |
| CC-3059 | SAMEA4731370 | ENA | (Ness et al. 2016) |
| CC-3063 | SAMEA4731374 | ENA | (Ness et al. 2016) |
| CC-3079 | SAMEA4731382 | ENA | (Ness et al. 2016) |
| CC-3084 | SAMEA4731383 | ENA | (Ness et al. 2016) |

Table S5: Oligo primers used or developed in this study.

| Taxa designed for | Name | Sequence (5'-->3') |
| --- | --- | --- |
| Eukaryotes (18S rRNA gene)* | TAReuk454FWD1 | CCAGCASCYGCGGTAATTCC |
| Eukaryotes (18S rRNA gene)* | TAReuk454REV3 | ACTTTCGTTCTTGATYRA |
| Prokaryotes (16S rRNA gene)** | 341F | CCTAYGGGRBGCASCAG |
| Prokaryotes (16S rRNA gene)** | 806R | GGACTACNNGGGTATCTAAT |
| Diatoms*** | rbcL-F | ATGTCTCAATCTGTAWCAGAACGGACTC |
| *Skeletonema marinoi/subsalsum* | rbcL-R | GATACCTGTAGCAGGACCTTGG |
| *Skeletonema marinoi* | Sm_C2W24-F | CATGAAACGGAAACTCTGAGCAC |
| *Skeletonema marinoi* | Sm_C2W24-R | AATATGCTGCGTTTCGAGTTCAATG |
| *Skeletonema marinoi* | Sm_C12W1-F | AGGYTTCGCCTCCTCAAAC |
| *Skeletonema marinoi* | Sm_C12W1-R | GGCACGATGCACACGCAAAG |
| *Skeletonema marinoi* | Sm_C12W2-F | CCCTCCAATACGCAACAATCC |
| *Skeletonema marinoi* | Sm_C12W2-R | CAATTCCACACGTCCCACC |
| *Skeletonema marinoi* | Sm_C16W4-F | CTGGTCCYATCAAGTGTGTGTCATC |
| *Skeletonema marinoi* | Sm_C16W4-R | TTCCCGTGAGGAGAGTGGAWG |
| *Chlorophytes* | Chlamy18SF | TGCCCTATCAACTTTCGATGGT |
| *Chlorophytes* | Chlamy18SR | GTGTGTACAAAGGGCAGGGA |
| *Chlamydomonas reinhardtii (mt-)* | Cr_Chr9W12-F | GTCACACTGCCCTGGTC |
| *Chlamydomonas reinhardtii(mt-)* | Cr_Chr9W12-R | CACCCACCTGTGCGATA |
| *Chlamydomonas reinhardtii(mt+)* | Cr_Chr1W3-F | TCGCTGCCACACTTCAC |
| *Chlamydomonas reinhardtii(mt+)* | Cr_Chr1W3-R | GGTTGGTCCAGCAGTCAG |
| *Chlamydomonas reinhardtii(mt+)* | Cr_Chr3W8-F | CTCCACATGGCCCTTAG |
| *Chlamydomonas reinhardtii(mt+)* | Cr_Chr3W8-R | CTGGTGAACCGGAATGT |
| *Chlamydomonas reinhardtii(mt+)* | Cr_Chr3W10-F | CAACGCGGCTGTTGGTC |
| *Chlamydomonas reinhardtii(mt+)* | Cr_Chr3W10-R | CGATCGGAGGTTTATATGCGTTTC |
| *-* | *Illumina F adapter* | ACACTCTTTCCCTACACGACGCTCTTCCGATCT |
| *-* | *Illumina R adapter* | GTGACTGGAGTTCAGACGTGTGCTCTTCCGATCT |

*(Stoeck et al. 2010), ** (Sundberg et al. 2013), ***(Guo et al. 2015)

Table S6. Optimization of denoising pipeline of amplicon sequencing data based on experimental observations.

Additional detail on each iteration is given in the *Attempted optimization of DADA2 parameters* section of the Supplemental Information.

| **Iteration** | Total ASVs^1^ | False negative ASVs^2^ | False positive ASVs^3^ | Read proportion TP^4^ | Read proportion FP^5^ | Mismatches between populations^6^ |
| --- | --- | --- | --- | --- | --- | --- |
| Exact matches (BBMerge without denoising) | 228,804 | 0 | 148,753 | 11.11% | 7.17% | 215 |
| ampliseq (integrate DADA2+Qiime2) | 1,006 | 110 | 0 | 0 | 0 | n/a |
| DADA2 with merging (DADA2) | 4,096 | 1 | 985 | 56.93% | 7.39% | 70 |
| Stringent DADA2 on amplicons (BBMerge+DADA2) | 1,062 | 3 | 16 | 17.89% | 0.03% | 1 |
| Relaxed DADA2 on amplicons (BBMerge+DADA2) | 1,483 | 0 | 346 | 17.81% | 1.19% | 4 |
| ^1^ – Total ASVs: the number of unique sequences produced from merged reads, either without (*Exact matches*) or with denoising.  ^2^ – False negative ASVs: the number of expected alleles that are missing from the total ASV count.  ^3^ – False positive ASVs: the number of unexpected alleles that are within 12bp of the expected 484bp barcode length, and contain the 5’ and 3’ regions identical among the expected alleles.  ^4^ – Read proportion TP: the percentage of the original read pairs that both merge successfully, and match one of the 110 true alleles.  ^5^ – Read proportion FP: the percentage of the original read pairs that both merge successfully, and match one of the identified false positive alleles.  ^6^ – Mismatches between populations: the number of times an allele expected exclusively in one population is seen in the other population where it was not actually present. In the data, there are 2,149 occasions where this would happen if all alleles were seen in all samples. | | | | | | |

Figure S1: PCR test of species specificity in amplification of the four *S. marinoi* barcoding loci and two *C. reinhardtii,* using DNA from other phytoplankton. All four *S. marinoi* loci were assayed in the close relative *Skeletonema subsalsum*, showing that only *Sm_C12W2* amplifies efficiently in this species (top panel). The second PCR (bottom panel) was performed only for *Sm_C12W1* but on a battery of microalgal samples that were mixed at equal *in vivo* chlorophyll *a* fluorescence density before DNA extraction. ‘Mix diatom’ corresponds to DNA from a mixture of seven diatom species isolated from the same Baltic Sea location as the *S. marinoi* strains (GP) and includes two different *Chaetoceros sp.* and *Fragilariopsis sp.*, and one each of *Thalassiosira baltica*, *Melosira sp.*, and *Cyclotella sp. ‘*Mixture’ corresponds to DNA from a mixture of the cyanobacteria *Nodularia spumigena*, *Aphanizomenon klebahnii*, *Dolichospermum sp.*, the dinoflagellate *Prorocentrum cordatum*, and a small (2-4 µm diameter) coccoid chlorophyte of unknown genus. ‘S_marinoi’ corresponds to DNA from strain VG1-2_86 which served as positive control for amplification of *Sm_C12W1*, and ‘NT’ is No DNA Template and negative controls. Panel C shows a lack of PCR amplification of two *Chlamydomonas reinhardtii* Barcode loci in two related species, *Chlamydomona* sp, and *Microglena* sp. Primer sequences are shown in Table S5.

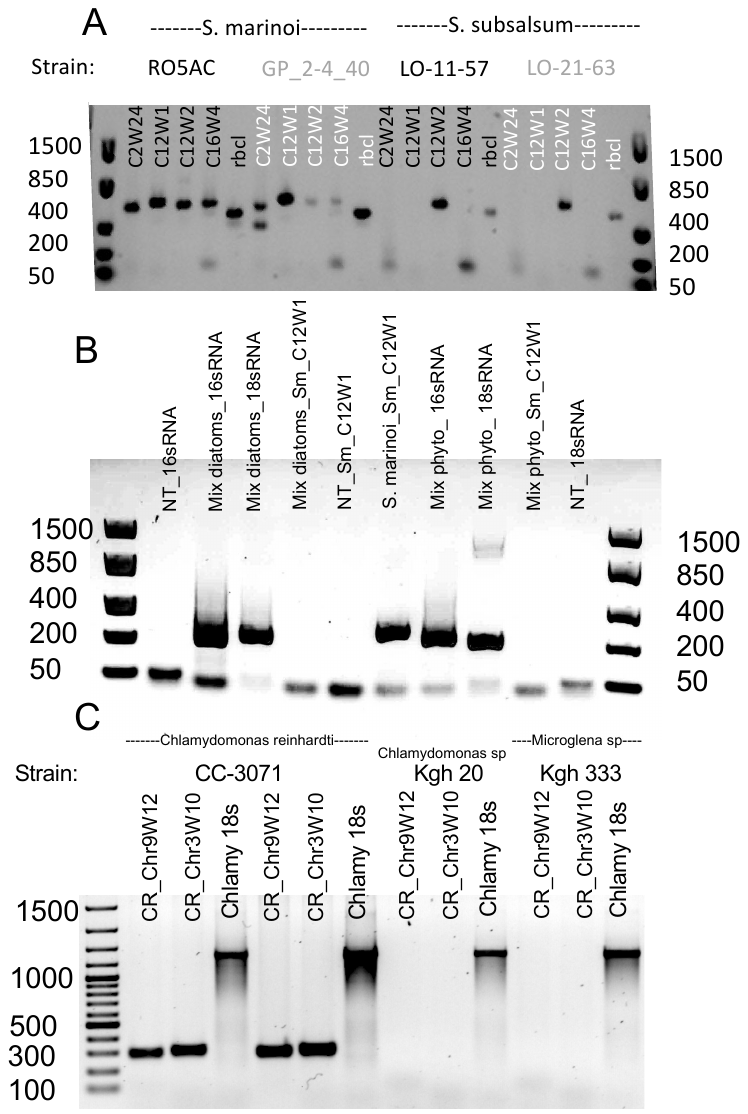

Figure S2: Pie charts depicting the breakdown of read pairs in each of the *S. marinoi* barcodes, based on the single-strain samples. Blue segments correspond to read pairs that did not merge; red segments correspond to read pairs that merged and matched the expected alleles for the given strain; and green segments correspond to unknowns, with a further breakdown of their identity in the outer ring.

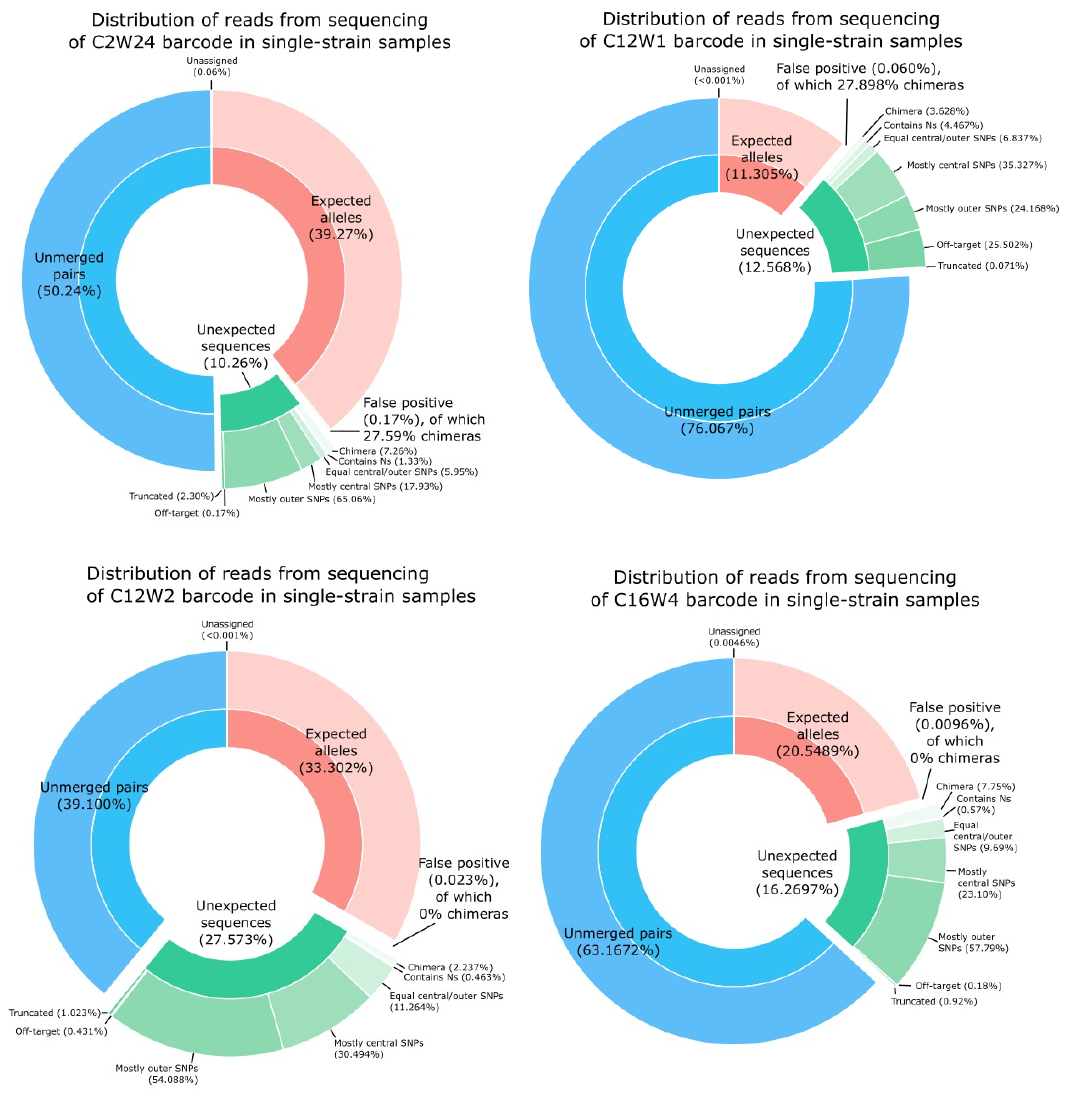

Figure S3: Mean error distribution across the length of the *Sm_C12W1* reads. Values are presented as the proportion of all reads labelled as either containing SNPs or as potential chimeras (using the Usearch uchime2_ref method as described in the Materials and Methods). Red positions denote SNP bases that differ between alleles in the database (i.e., those that are informative as to the strain’s identity, but also the ones that will be flagged as errors from PCR chimeras). Panel A depicts the result when the reads are mapped to a consensus sequence of the strain’s alleles (i.e. masking the influence of the chimeras). Panel B depicts the result when the reads are mapped to a reference containing the individual alleles.

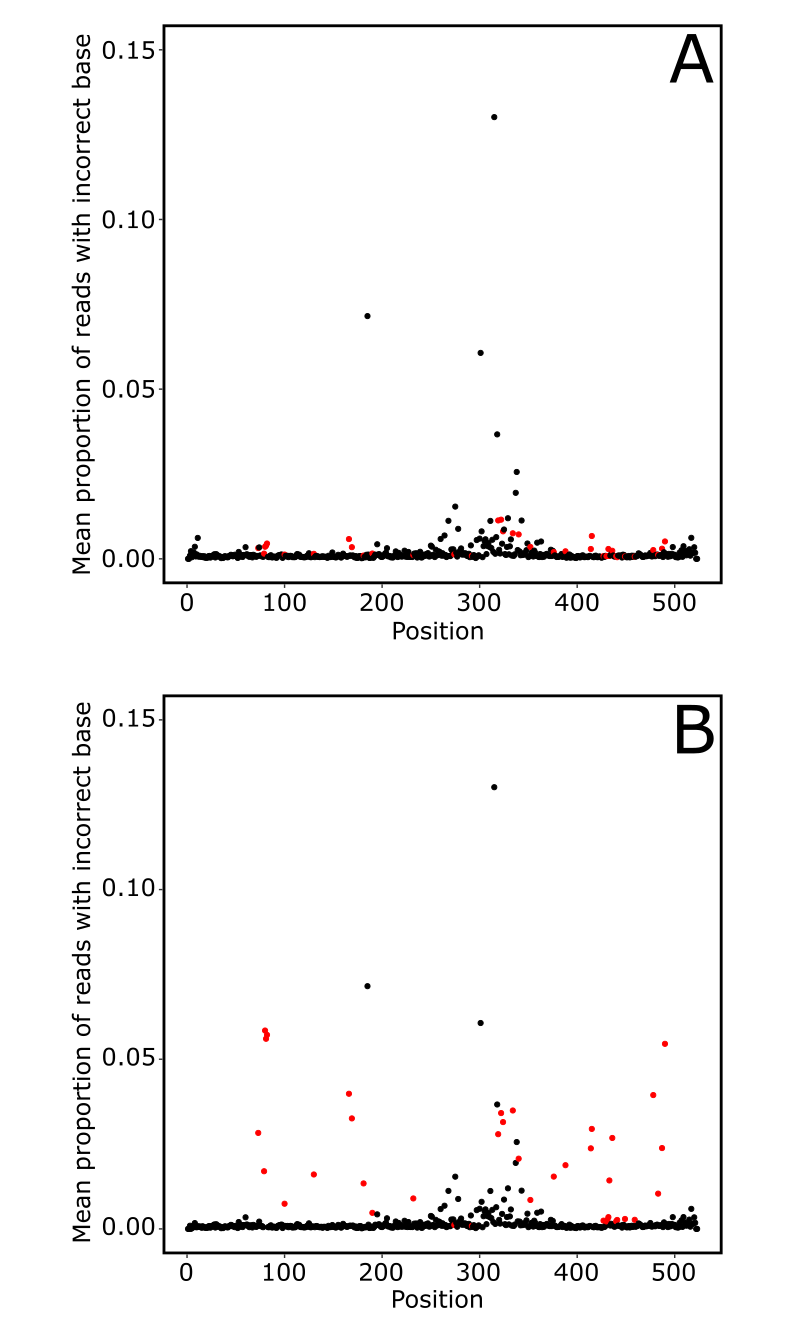

Figure S4: Allele correlations for 58 strains of *S. marinoi* across the 42-day selection experiment based on exact sequence matches and without de-noising by DADA2. Shared homologous alleles are indicated by alphabetical indexes in the corner of each allele (i.e., lower right equals allele one, upper left allele two), and their abundances have been allocated between strains using differential equations. All such equations assume a 1:1 ratio between alleles, except allele two in GP2-4_43, where a ratio of 0.15 is used based on the genotype sample observation. The black line across panels corresponds to the expected 1:1 ratio between heterozygous alleles. The blue line is a second order polynomial function fitted to the data, with shaded area corresponding to 95% confidence interval. Strain GP2-4_32, Strain VG1-2_65or99 and VG1-2_99or65's alleles, were not sequenced but manually identified in an iterative process from the pool ASV that DADA2 predicted for the artificial evolution experiments.
